# Activation of the proton-sensing GPCR, GPR65 on fibroblast-like synoviocytes contributes to inflammatory joint pain

**DOI:** 10.1101/2024.04.29.590277

**Authors:** Luke A. Pattison, Rebecca H. Rickman, Helen Hilton, Susanne N. Wijesinghe, Graham Ladds, Li Yang, Simon W. Jones, Ewan St. John Smith

**Author notes:** To whom correspondence should be addressed: Luke A. Pattison: Department of Pharmacology, University of Cambridge;* *or Ewan St. John Smith: Department of Pharmacology, University of Cambridge, UK;.

## Abstract

Inflammation is associated with localised acidosis, however, attributing physiological and pathological roles to proton-sensitive receptors is challenging due to their diversity and widespread expression. Here, agonists of the proton-sensing GPCR, GPR65, were systematically characterised. The synthetic agonist BTB09089 (BTB) recapitulated many proton-induced signalling events and demonstrated selectivity for GPR65. BTB was used to show that GPR65 activation on fibroblast-like synoviocytes (FLS), cells that line synovial joints, results in the secretion of pro-inflammatory mediators capable of recruiting immune cells and sensitising sensory neurons. Intra-articular injection of BTB resulted in GPR65-dependent sensitisation of knee-innervating neurons and nocifensive behaviours in mice. Stimulation of GPR65 on human FLS also triggered the release of inflammatory mediators and synovial fluid samples from human osteoarthritis patients were shown to activate GPR65. These results suggest a role of GPR65 in mediating cell-cell interactions that drive inflammatory joint pain in both mice and humans.

## Introduction

Chronic inflammatory conditions, such as arthritis and inflammatory bowel disease (IBD) are associated with pain, a leading complaint of those living with such diagnoses, and poorly managed by current therapeutics^1–3^. As well as pain, inflammatory conditions often coincide with localised acidosis^4,5^, owing to immune cell influx^6^, tissue damage^7^ and regulated release of protons in response to inflammatory stimuli^8^. The resulting decrease in extracellular pH exacerbates inflammation: under acidic conditions immune cells produce and secrete more pro-inflammatory mediators^9^; additionally, protons may directly activate and sensitise sensory neurons^10,11^. Furthermore, subdermal application of acidic solutions in humans can cause pain independent of inflammation^12,13^. These findings suggest that inflammatory acidosis may contribute to the pain experienced by patients. Therefore, receptors sensitive to extracellular pH might represent novel points of therapeutic intervention for treating inflammatory pain.

Several receptor families are tuned to detect changes in extracellular pH, including: acid-sensing ion channels, certain transient receptor potential channels and two-pore potassium channels, and proton-sensing G protein-coupled receptors (PS-GPCRs). While the contributions of acid-sensitive ion channels to inflammatory pain are relatively well understood, the involvement of the six PS-GPCRs is less clear^14^. PS-GPCRs are expressed by several cell types involved in inflammation, including immune cells^15^, fibroblasts^16^, and sensory neurons^17^, with increased expression reported for pre-clinical models of inflammation, as well as human pathologies^18–24^.

Among the PS-GPCRs, GPR65 (also referred to as T Cell Death Associated Gene 8, TDAG8), shows the greatest upregulation in inflammation^18,25^. Furthermore, GPR65 knockout (KO) mice exhibit reduced pain behaviours following localised injection of acid^26^, or induction of experimental arthritis^27^, which has been linked to reduced infiltration of immune cells^28^. Localised GPR65 knockdown also reduces the severity of mechanical hypersensitivity evoked by inflammatory insult, localised injection of acid or a GPR65 agonist^29^. GPR65 has also been linked to the pathology of visceral pain conditions, including IBD^30^ and cirrhosis^31^. Genetic associations of *GPR65* and human pathologies including chronic obstructive pulmonary disorder^32^, ankylosing spondylitis^33^, ulcerative colitis^23^ and atopic dermatitis^34^ have also been described. Thus, understanding the signalling mechanisms of GPR65 could be of benefit across a broad range of conditions, making it an ideal candidate for elucidating the potential role of PS-GPCRs in inflammatory pain.

The widespread expression of proton-sensitive receptors makes attributing contributions of individual receptors to physiological processes and pathology a challenging task. However, in addition to protons^35^, GPR65 is reported to be activated by the glycosphingolipid, psychosine^36^, and the synthetic agonist BTB09089 (BTB)^37^. This study firstly characterises GPR65 signalling in a recombinant system, and confirms selectivity of BTB as a GPR65 agonist, before leveraging that selectivity to determine the consequences of GPR65 activation in more physiologically relevant cell types (sensory neurons and fibroblasts), and demonstrating the role of GPR65 in mediating cell-cell interactions that underpin inflammatory joint pain in mice and humans.

## Results

### BTB recapitulates a similar intracellular signalling signature to protons at GPR65

GPCRs coordinate pleiotropic intracellular signalling events, further complexity being afforded by functional selectivity, also known as biased signalling: the ability of distinct agonists to drive different responses through the same receptor. Exploring signalling bias at GPR65 among protons, BTB and psychosine could thus represent a valuable opportunity to bypass the promiscuity of proton-elicited activation in primary cells. This is because if either BTB or psychosine recapitulate the signalling signature of protons following GPR65 activation, they are more likely to be selective for GPR65 over other proton-sensitive receptors. Therefore, the ability of each agonist to evoke signalling responses was assayed in a uniform cellular background, expressing mouse GPR65 as the only known proton-sensitive receptor (mGPR65-CHO cells; Fig. 1a). Consistent with previous reports^35^, increasing the proton concentration resulted in cAMP accumulation until pH < 6.4, when production decreased (Fig. 1b). Importantly proton-stimulation of parental Flp-IN cells caused no cAMP accumulation (F(1,17) = 0.139, *p =* 0.713; Supplementary Fig. 1a) indicating the dependence of proton-induced cAMP accumulation on GPR65 expression. Like protons, BTB also caused a concentration-dependent increase in cAMP accumulation in mGPR65-CHO cells (Fig. 1b). Psychosine is reported to decrease forskolin-induced cAMP accumulation^36^, cells exposed to psychosine were therefore co-stimulated with 1 µM forskolin, and, as expected a concentration-dependent decrease in cAMP production was observed (Fig. 1b). Next, the ability of protons, BTB and psychosine to stimulate Ca^2+^ release from intracellular stores was assessed using the Ca^2+^-sensitive dye Fluo4, only psychosine and protons evoked Ca^2+^ mobilisation at concentrations above 1 µM and pH < 6.6, respectively (Fig. 1c). To confirm the observed increases in [Ca^2+^] were due to engagement of GPR65, experiments were repeated using parental Flp-IN cells. 10 µM psychosine having no effect (10 µM psychosine: Flp-IN, 0.33 ± 1.62 %, mGPR65-CHO, 85.37 ± 6.86 %, *p-adj* = 0.007; Supplementary Fig. 1b).

**Figure 1.**
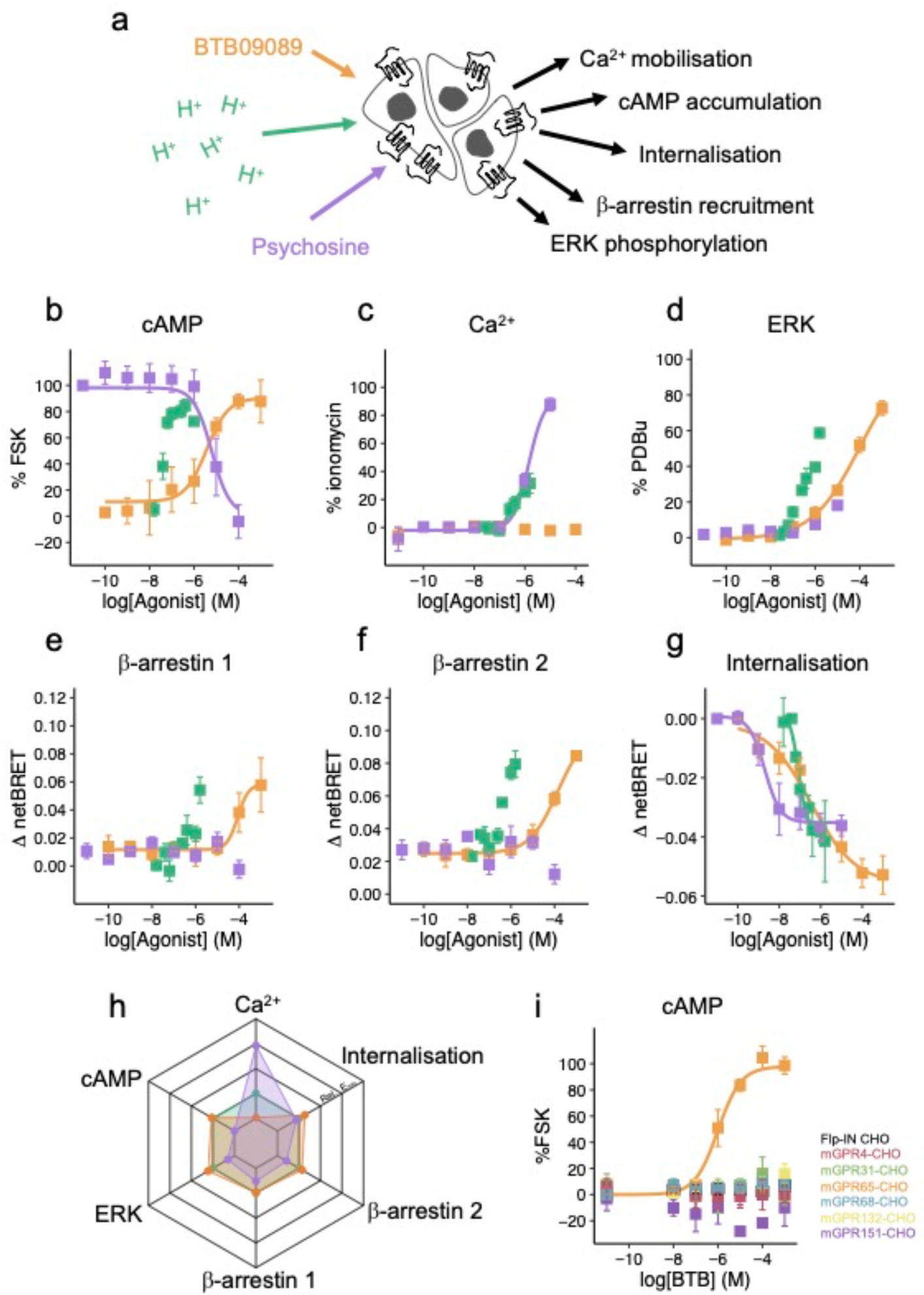
BTB recapitulates a similar intracellular signalling signature to protons at GPR65. **(a)** Intracellular signalling responses coordinated by mouse GPR65 were assessed in a CHO cell background, the ability of protons (green), BTB (orange) or psychosine (purple) to coordinate **(b)** accumulation of cAMP (FSK, forskolin), **(c)** intracellular Ca^2+^ mobilisation, **(d)** phosphorylation of ERK1/2 (PDBu, phorbol 12,13-dibutyrate), recruitment of **(e)** β-arrestin 1 or **(f)** β-arrestin 2 and **(g)** receptor internalisation was assessed. **(h)** To compare the signalling profiles of each GPR65 agonist, the peak response of each agonist in each pathway was normalised to that achieved by proton stimulation. The E_max_ of psychosine in the cAMP pathway was set as 0. **(h)** The selectivity of BTB among other PS-GPCRs was assessed using stable cell lines and the cAMP assay. Data are from at least three independent experiments where each [agonist] was assayed in duplicate.

Although no response was seen at pH 7, a small response was recorded when Flp-IN cells were exposed to pH 6, but this was lower than that seen in mGPR65-CHO cells (pH 6: Flip-IN, 7.58 ± 2.20 %, mGPR65-CHO, 25.36 ± 0.17 %, *p-adj* = 0.006; Supplementary Fig. 1b), indicating that protons can induce a GPR65-dependent mobilisation of intracellular Ca^2+^, but that Fluo4 may have some inherent sensitivity to high concentrations of protons. Another facet of GPCR signalling is ERK activation: all three agonists coordinated ERK1/2 activation, however, efficacy differed with BTB producing the greatest response, followed by protons and lastly psychosine (Fig. 1d). Protons could not activate ERK1/2 in parental Flp-IN cells, thus highlighting the necessity of GPR65 for the observed responses in mGPR65-CHO cells (pH 6: Flp-IN, 1.11 ± 0.15 %, mGPR65-CHO, 39.51 ± 2.16 %, t = -17.756, df = 2.018, *p =* 0.003; Supplementary Fig. 1c). β-arrestins mediate GPCR desensitisation of GPCRs, but can also coordinate signalling events upon recruitment to active receptors. GPR65-mediated β-arrestin recruitment was assessed using a BRET-based assay to quantify the proximity of YFP-tagged β-arrestins to a luciferase tagged GPR65 (mGPR65-RLuc8). Following 15 minutes of exposure to protons or BTB, increased BRET was detected between GPR65-RLuc8 and ý-arrestin 1-YFP (Fig. 1e). No change in BRET was observed upon psychosine treatment, suggesting that it does not coordinate β-arrestin recruitment to GPR65. Similar results were obtained for β-arrestin 2, with only protons and BTB able to evoke recruitment (Fig. 1f). Following activation many GPCRs are internalised. To assess agonist-induced GPR65 internalisation, BRET assays to quantify the proximity of mGPR65-RLuc8 to a fluorescent fusion protein resident in the plasma membrane (RIT-venus) were conducted. Following stimulation with protons, BRET between mGPR65-RLuc8 and RIT-venus decreased in a concentration dependent manner; BTB and psychosine also induced receptor internalisation (Fig. 1g).

For a more comprehensive comparison of GPR65 signalling, the maximum effect (E_max_) of each agonist for each output was normalised to that achieved by proton-stimulation. This revealed that BTB has a very similar signalling fingerprint to protons, only differing in its inability to mobilise intracellular Ca^2+^, whereas psychosine coordinates a very different response profile in mGPR65-CHO cells (Fig. 1h). Thus, BTB appears to represent a useful tool to mimic proton-induced signalling. A full assessment of signalling bias was not appropriate given the discrepancies in the range of concentrations tested for each agonist. Finally, the selectivity of BTB was confirmed against other PS-GPCRs using the cAMP accumulation assay. BTB only coordinated a response in cells expressing mGPR65 (Fig. 1i) and thus represents a selective tool for stimulating a similar intracellular signalling cascade as protons via GPR65, therefore enabling its use to study GPR65 driven physiology in cells and systems that express multiple proton-sensitive receptors.

### Intra-articular injection of BTB causes inflammation and pain-like behaviours in mice

Having established that BTB recapitulates most features of proton-induced GPR65 signalling and is selective for GPR65 over other PS-GPCRs, the contribution of GPR65 to inflammatory joint pain was investigated. Intra-articular BTB injection served to selectively activate GPR65 in the joint environment, mimicking the increased acidity of synovial fluid common to human arthritis^38–41^. Given the higher prevalence of arthritis in females^42,43^, initial studies were restricted to female mice. After capturing baseline behaviours, mice received a unilateral intra-articular injection of either 100 µM BTB or DMSO (0.1% v/v, vehicle control), and were studied for 7 days (Fig. 2a). Swelling of the injected joint, as indicated by increased ipsilateral vs. contralateral knee width was observed, dependent on both the substance injected and time (interaction: injection:time: F(7,112) = 8.320, *p* < 0.0001; Fig. 2b). Peak inflammation occurred 24-hours post-injection of BTB (knee width ratio at 24-hours: BTB, 1.19 ± 0.02, DMSO, 1.00 ± 0.01, t = 9.376, df = 7, *p-adj* < 0.0001; Fig. 2b), after which swelling subsided during the experimental period. No change in knee width was observed following intra-articular DMSO injection (F(1,124) = 0.003, *p-adj* = 0.955; Fig. 2b). In line with knee inflammation, increased pain-like behaviours were also recorded dependent on the substance injected, including increased mechanical hypersensitivity of the BTB injected knee (F(1,84) = 34.34, *p* < 0.0001; Fig. 2c), an evoked pain response. The mechanical sensitivity of the contralateral joint was unaffected (F(1,84) = 0.801, *p =* 0.373; Supplementary Fig. 2a). The digging behaviour of mice, an ethological readout of animal pain and wellbeing, was also negatively affected by BTB injection, with increased latency to dig (F(1,84) = 25.40, *p* < 0.0001; Fig. 2d) and reductions in both digging duration (F(1,84) = 15.95, *p* = 0.00014; Fig. 2e) and number of burrows produced (F(1,84) = 45.59, *p* > 0.0001; Fig. 2f) observed. These findings likely mimic the withdrawal from everyday activities reported by human pain patients, as a result of spontaneous pain and apathy experienced as part of their inflammation^3^, given mice retained full ability to use the injected joint, as inferred by the lack of an effect of injections on performance in the rotarod test (F(1,84) = 0.351, *p* = 0.555; Supplementary Fig. 2b). The swelling and painlike behaviours observed following BTB injection into the mouse knee joint suggest that GPR65 activation is sufficient to drive inflammatory pain. The next objective was to determine the cellular basis of GPR65-mediated inflammatory pain.

**Figure 2.**
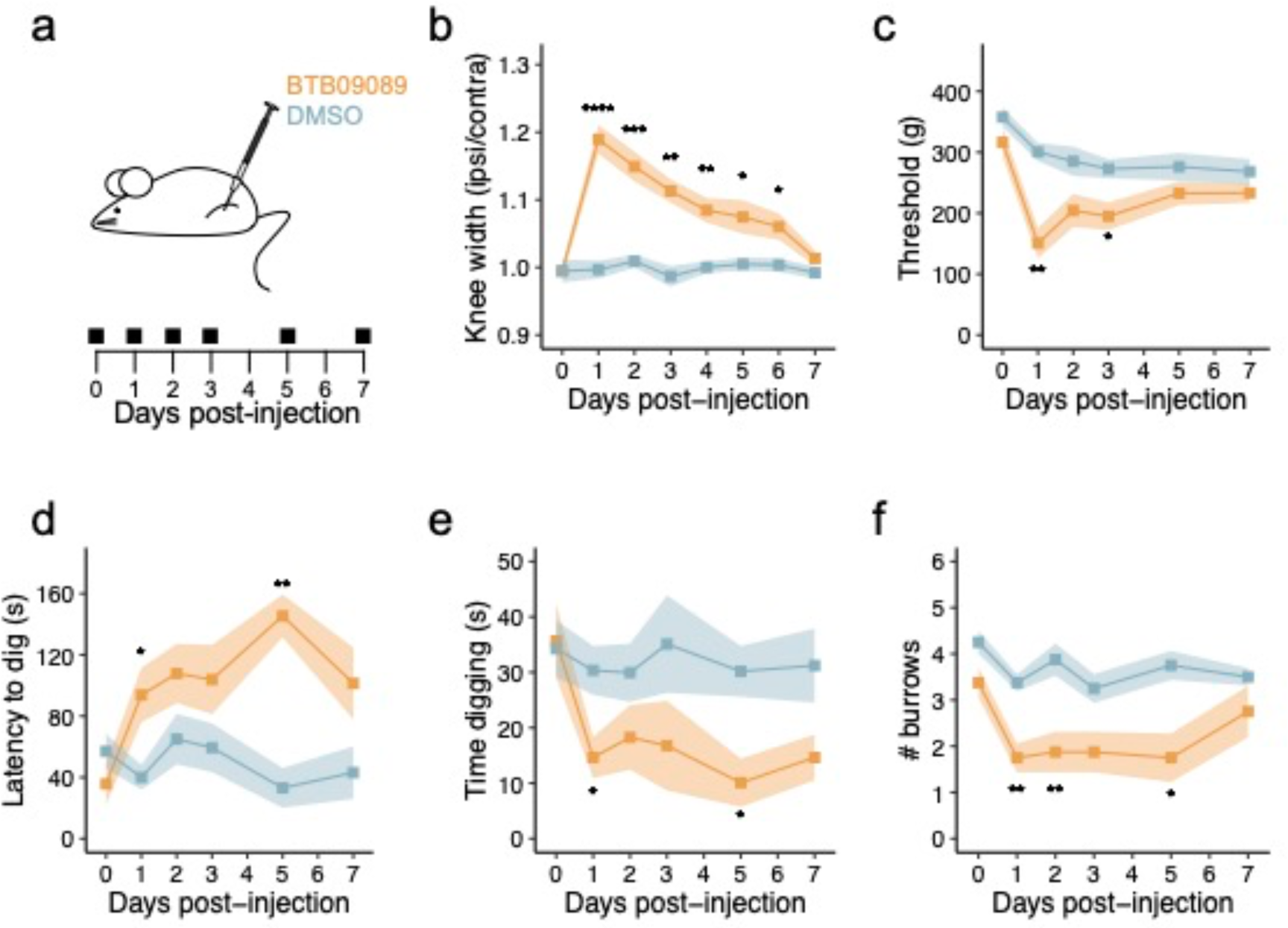
Intra-articular injection of BTB causes inflammation and pain-like behaviours in mice. **(a)** Schematic representation and experimental timeline. Female mice received a unilateral injection of either 100 µM BTB (orange) or 0.1% (v/v) DMSO (blue). **(b)** The ratio of the ipsilateral to contralateral knee width was calculated as a measure of the extent of inflammation. **(c)** Mechanical sensitivity of injected knee joints was determined by pressure application measurement. The **(d)** latency to dig, **(e)** time spent digging and **(f)** number of burrows dug were also measured across experimental time. * *p-adj* < 0.05, ** *p-adj* < 0.01, *** *p-adj* < 0.001, **** *p-adj* < 0.0001: Repeated-measures analysis of variance followed by Bonferroni-corrected post hoc; annotated statistical differences are the effect of injected substance for each day of testing. N = 8 female mice per group.

### BTB-induced sensitisation of sensory neurons depends upon cells resident in the joint

Sensory neurons, whose cell bodies reside in the dorsal root ganglia (DRG), are the principal orchestrators of nociception, transmitting noxious stimuli to the brain where it manifests as pain^44^. Under inflammatory conditions sensory neurons are sensitised by mediators released by numerous cell types, giving rise to hyperalgesia, i.e. a gain in pain^45^. Given the pain-like behaviours observed following BTB injection, it was hypothesised that BTB activates GPR65 on nociceptors, triggering intracellular signalling events, that sensitise these cells, leading to increased transmission of noxious stimuli and increased pain. To assess this, naïve mouse sensory neurons were dissociated and incubated with BTB, DMSO or regular media overnight (Fig. 3a), akin to the greatest pain responses being seen 24-hours post-intra-articular injection (Fig. 2). Electrophysiological characterisation was then performed to determine cellular excitability, by investigating the amount of current required to evoke action potential discharge (rheobase; Fig. 3b). However, there was no difference in rheobase across the three conditions (Media, 372.41 ± 45.33 pA, DMSO, 274.67 ± 52.42 pA, BTB, 331.48 ± 59.9 pA; F(2,83) = 0.898, *p =* 0.411; Fig. 3b-c). Similarly, no effect of any culturing condition on other intrinsic or active DRG neuron properties was measured (Table 1).

**Figure 3.**
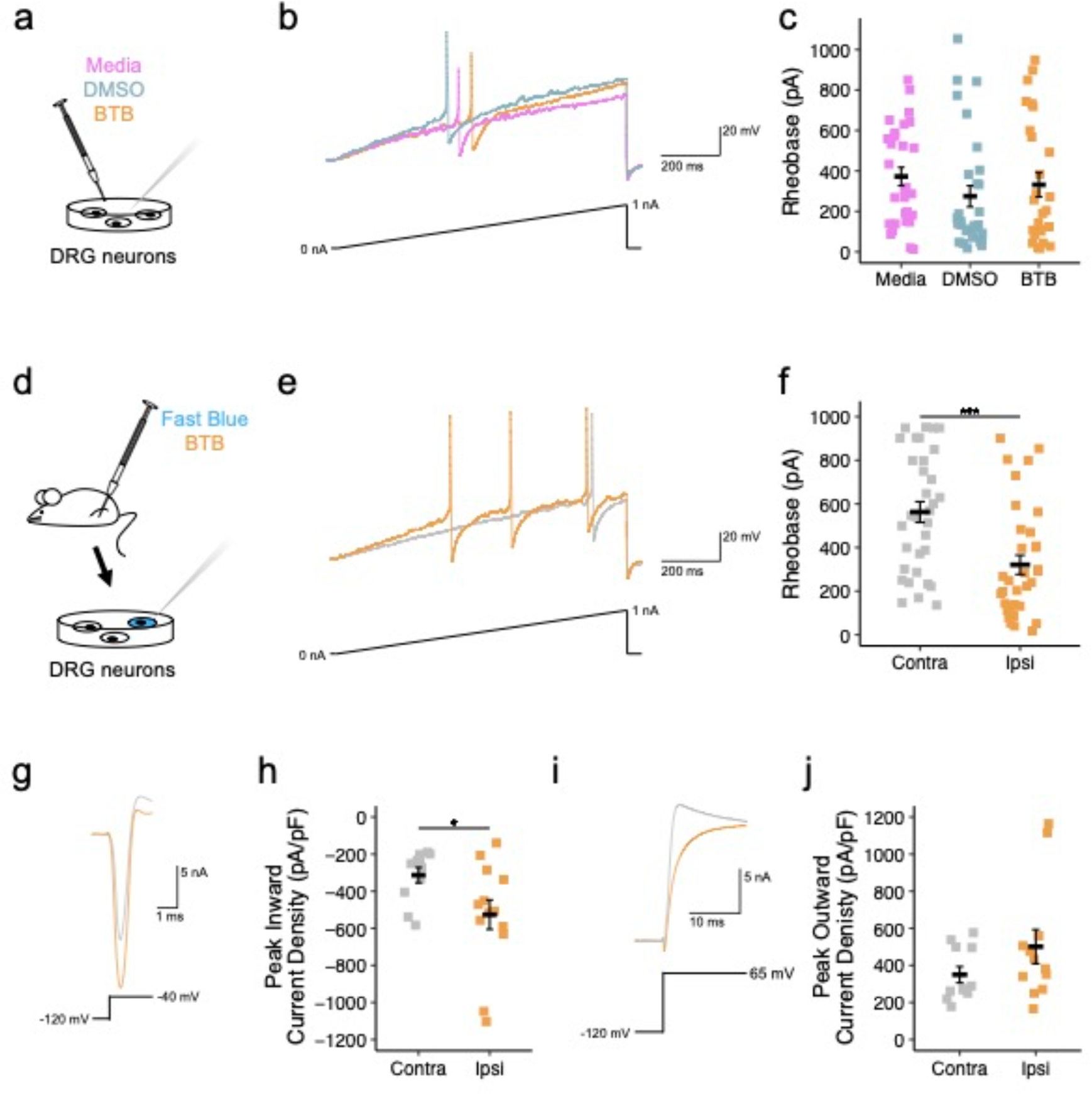
BTB-induced sensitisation of sensory neurons depends upon cells resident in the joint. **(a)** Naïve DRG neurons were cultured overnight with 100 µM BTB (orange), 0.1% (v/v) DMSO (blue) or regular culture media (pink), before electrophysiological characterisation. **(b)** Representative current clamp recordings of neurons of comparable capacitance, showing action potentials evoked by ramp injection of current (0–1 nA, 1 s). **(c)** Stepwise current injections were used to determine the rheobase of sensory neurons. **(d)** Following retrograde labelling of knee-innervating sensory neurons (with Fast Blue), mice were injected with 100 µM BTB into one knee, 24-hours post-injection DRG from the ipsilateral (Ipsi, orange) and contralateral (Contra, grey) were collected and cultured for electrophysiological characterisation. **(e)** Representative current clamp recordings of neurons of comparable capacitance, showing action potentials evoked by ramp injection of current (0–1 nA, 1 s). **(f)** Stepwise current injections were used to determine the rheobase of sensory neurons. **(g)** Representative inward current trace from current-voltage (IV) analyses, elicited by a -40 mV test pulse in cells of comparable capacitance. **(h)** Peak inward current density from IV analyses. **(i)** Representative outward current trace from IV analyses, elicited by a 65 mV test pulse in cells of comparable capacitance. **(j)** Peak outward current density from IV analyses. * *p* < 0.05, *** *p* < 0.001: **(c)** One-way ANOVA followed by Bonferroni-corrected post-hoc, **(f,h,j)** unpaired t-test.

**Table 1:** Intrinsic and active properties of naïve dorsal root ganglion neurons cultured in regular media, DMSO or BTB overnight.

|  | **Media (n = 29)** | **DMSO (n = 30)** | **BTB (n = 27)** |
| --- | --- | --- | --- |
| Resting membrane potential (mV) | -43.79 ± 0.83 | -46.13 ± 1.00 | -44.82 ± 0.92 |
| Capacitance (pF) | 45.40 ± 4.31 | 39.65 ± 4.52 | 42.58 ± 4.78 |
| # APs at suprathreshold | 1.80 ± 0.70 (n = 10) | 3.00 ± 0.47 (n = 23) | 3.71 ± 0.72 (n = 14) |
| Action potential amplitude (mV) | 75.62 ± 5.20 | 82.35 ± 3.44 | 78.49 ± 3.85 |
| Half peak duration (ms) | 2.11 ± 0.28 | 2.64 ± 0.27 | 2.26 ± 0.25 |
| Afterhyperpolarisation amplitude (mV) | 15.85 ± 0.87 | 14.94 ± 0.92 | 15.95 ± 0.83 |
| Afterhyperpolarisation duration (ms) | 16.84 ± 2.66 | 22.71 ± 4.48 | 15.11 ± 2.42 |

Considering that mice injected with BTB exhibited pain-like behaviours (Fig. 2), studies were next restricted to the knee-innervating neurons, identified by retrograde tracing, of mice that had received an intra-articular BTB injection (Fig. 3d). As before, this cohort exhibited knee inflammation 24-hours post-injection, alongside increased mechanical sensitivity and a decrease in digging behaviour (Supplementary Fig. 3). Interrogation of fast blue positive, i.e. knee-innervating, neurons revealed higher excitability of cells which projected to the inflamed knee compared to those supplying the non-injected knee, as shown by their lower rheobase (Ipsi, 320.88 ± 44.00 pA vs Contra, 562.73 ± 47.62 pA, t = 3.73, df = 64.427, *p* = 0.0004; Fig. 3e-f). No difference was seen in any other parameter measured (Table 2). To better understand the cellular changes resulting in the greater excitability of neurons projecting to the inflamed joint, macroscopic voltage-gated currents were studied, a greater inward current density being measured for ipsilateral neurons, (Ipsi, -526.24 ± 79.14 pA/pF, Contra, -313.16 ± 41.65 pA/pF, t = 2.383, df = 17.919, *p* = 0.029; Fig. 3g-h); no difference was observed in outward current density (Ipsi, 566.62 ± 106.99 pA/pF, Contra, 350.04 ± 43.81 pA/pF, t = - 1.873, df = 15.828, *p* = 0.079; Fig. 3i-j). Considered together, these results demonstrate that exposure of sensory neurons to BTB alone is insufficient to increase their excitability and suggest that other cell types within the joint environment, are necessary for BTB to induce neuronal hyperexcitability and nocifensive behaviours.

**Table 2:** Intrinsic and active properties of knee-innervating dorsal root ganglion neurons from the BTB-injected side (Ipsi) and contralateral (Contra) side of mice. Erev, reversal potential; V1/2, half-activating potential of voltage-sensitive channels.

| **Current-clamp** | **Contra (n = 33)** | **Ipsi (n = 34)** |
| --- | --- | --- |
| Resting membrane potential (mV) | -46.09 ± 0.93 | -48.68 ± 1.16 |
| Capacitance (pF) | 53.96 ± 5.10 | 43.08 ± 5.23 |
| # APs at suprathreshold | 1.42 ± 0.23 (n = 12) | 1.86 ± 0.42 (n = 21) |
| Action potential amplitude (mV) | 80.76 ± 4.26 | 81.13 ± 4.53 |
| Half peak duration (ms) | 2.41 ± 0.50 | 2.23 ± 0.23 |
| Afterhyperpolarisation amplitude (mV) | 17.56 ± 0.64 | 17.43 ± 0.62 |
| Afterhyperpolarisation duration (ms) | 10.40 ± 1.29 | 10.84 ± 1.85 |
| **Voltage-clamp** | **Contra (n = 11)** | **Ipsi (n = 13)** |
| Inward: Erev | 62.83 ± 8.92 | 61.78 ± 10.26 |
| Inward: V1/2 | -37.69 ± 3.00 | -44.26 ± 2.86 |

### Fibroblast-like synoviocytes express GPR65 and respond to BTB stimulation

The involvement of multiple cell types in inflammation is becoming ever more apparent, particularly for arthritic conditions, where chondrocytes, immune cells, and osteoclasts have all been implicated in disease progression^46–48^. Fibroblast-like synoviocytes (FLS), which line the synovial joints and function to secrete synovial fluid, have also been implicated in inflammatory arthritis pathology and pain^49–51^ FLS also express GPR65^52,53^ and thus the consequences of engaging GPR65 on FLS were studied. Mouse FLS cells cultured from patellae express FLS markers, such as Cdh-11 and Cdh-248, as determined by immunoreactivity and qPCR (Fig. 4a-b), whereas expression of endothelial (Cd-31) and immune markers (Cd-68) was low (Fig. 4b), thus indicating high purity. qPCR analysis also revealed that among the PS-GPCRs, GPR65 was most highly expressed by FLS (Fig. 4b). In agreement with the expression data, stimulation of mouse FLS with either BTB or acidic pH induced cAMP accumulation (100 µM BTB, 3.72 ± 0.17 nM vs vehicle, 0.78 ± 0.17 nM, t = 9.24, df = 3, *p-adj* = 0.016; pH 6, 4.00 ± 0.16 nM vs vehicle, 0.78 ± 0.17 nM, t = 11.88, df = 3, *p-adj* = 0.008;Fig. 4c), consistent with the signalling responses seen in mGPR65-CHO cells (Fig. 1a).

**Figure 4.**
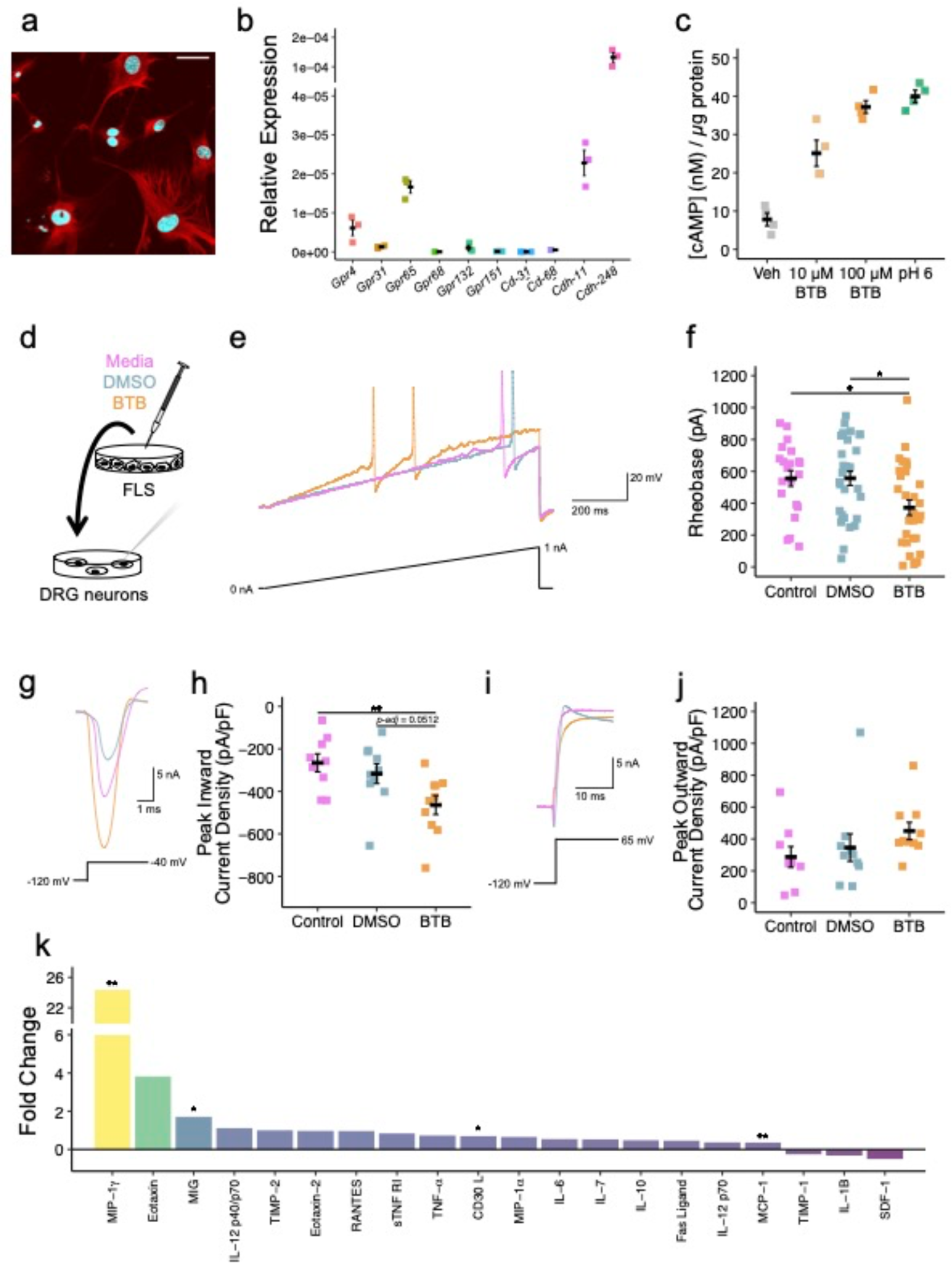
Fibroblast-like synoviocytes express GPR65 and respond to BTB stimulation. **(a)** Mouse fibroblast-like synoviocytes (FLS) express CDH-11 (Red: αCDH-11, Blue: Nuclear stain; Scale bar = 50 µm). **(b)** FLS gene expression was further interrogated via qPCR. **(c)** Intracellular FLS cAMP concentration following stimulation with pH 7.4 vehicle, BTB or pH 6 solution. **(d)** FLS were cultured overnight with either BTB (orange), DMSO (blue) or regular culture media (pink), 24-hours post-stimulation media were collected and later incubated with naïve DRG neurons overnight before electrophysiological characterisation. **(e)** Representative current clamp recordings of neurons of comparable capacitance, showing action potentials evoked by ramp injection of current (0–1 nA, 1 s). **(f)** Stepwise current injections were used to determine the rheobase of sensory neurons. **(g)** Representative inward current trace from IV analyses, elicited by a -40 mV test pulse in cells of comparable capacitance. **(h)** Peak inward current density from IV analyses. **(i)** Representative outward current trace from IV analyses, elicited by a 65 mV test pulse in cells of comparable capacitance. **(j)** Peak outward current density from IV analyses. **(k)** BTB-induced fold change in detection of inflammatory cytokines in conditioned media from stimulated FLS. * *p-adj* < 0.05, ** *p-adj* < 0.01: **(f, h, j)** One-way ANOVA followed by Bonferroni-corrected post-hoc. **(k)** Two-way ANOVA followed by Bonferroni-corrected post-hoc.

To ascertain whether FLS contribute to BTB-driven neuronal hyperexcitability and pain-like behaviour, FLS were stimulated with BTB or DMSO overnight before collection of media the following day. After dissociation of naïve mouse DRG, neurons were cultured overnight in the conditioned media collected from stimulated FLS prior to electrophysiological characterisation (Fig. 4d). A lower rheobase of neurons incubated in the media taken from BTB-stimulated FLS compared to media from either DMSO-treated FLS or unstimulated FLS cultures was observed (Media, 555.24 ± 49.06 pA, DMSO, 555.67 ± 44.95 pA, BTB, 372.00 ± 48.09 pA; F(2,78) = 5.22, *p* = 0.007; Fig. 4e-f), akin to the hyperexcitability previously seen in knee-innervating neurons isolated from BTB-injected mice (Fig. 3f). No difference was seen in any other parameter measured (Table 3). Analysis of macroscopic voltage-gated currents also revealed a larger inward current density of neurons cultured in media taken from BTB-stimulated FLS (Media, -265.84 ± 42.40 pA/pF, DMSO, -316.94 ± 46.01 pA/pF, BTB, -468.19 ± 40.55 pA/pF; F(2,27) = 6.115, *p* = 0.006; Fig. 4g-h), with outward currents mostly unaffected (Media, 287.14 ± 65.31 pA/pF, DMSO, 345.91 ± 86.20 pA/pF, BTB, 466.85 ± 51.85 pA/pF; F(2,27) = 1.811, *p* = 0.183; Fig. 4i-j).

**Table 3:**
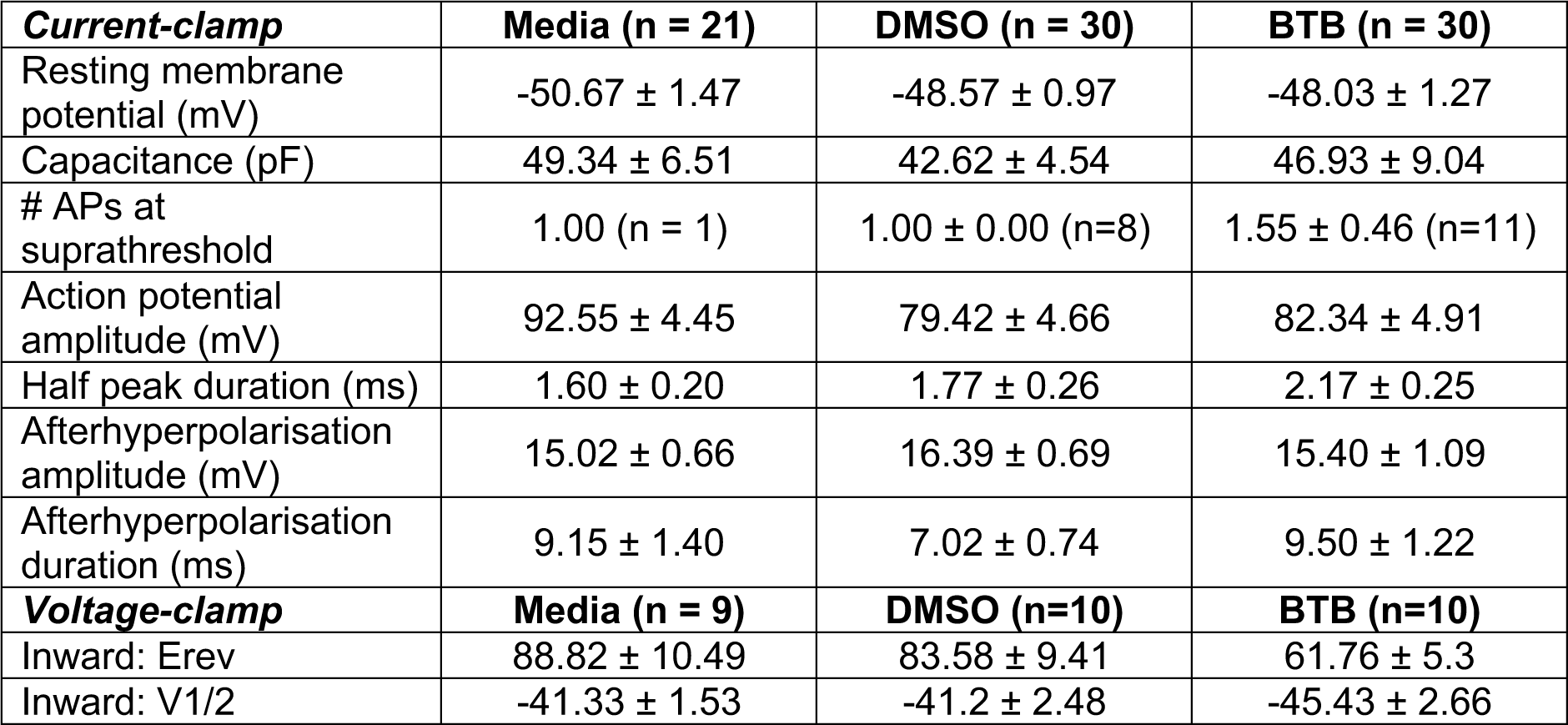
Intrinsic and active properties of naïve dorsal root ganglion neurons cultured in conditioned media collected from WT FLS stimulated with regular media, DMSO or BTB overnight.

Taken together these data suggest that BTB stimulation of FLS is able to re-create the hyperexcitability of sensory neurons taken directly from BTB-injected mice. To better understand the crosstalk between BTB-activated FLS and sensory neurons, the levels of inflammatory cytokines present in media taken from BTB-stimulated FLS were examined. Compared to DMSO-stimulation, exposure of FLS to BTB resulted in increased levels of many inflammatory cytokines, most notably macrophage inflammatory protein-1γ (MIP-1γ; BTB-stimulated density, 2.19 ± 0.22, DMSO-stimulated density, 0.09 ± 0.01, *p-adj =* 0.0044; Fig. 4k), monokine induced by γ interferon (MIG; BTB, 0.10 ± 0.003, DMSO, 0.04 ± 0.01, *p-adj =* 0.039; Fig. 4k), CD30 ligand (BTB, 0.21 ± 0.01, DMSO, 0.12 ± 0.02, *p-adj =* 0.043; Fig. 4k) and monocyte chemoattractant protein 1 (MCP1; BTB, 1.44 ± 0.03, DMSO, 1.05 ± 0.01, *p-adj =* 0.007; Fig. 4k; full analysis of cytokine levels are detailed in Table S1). These 4 cytokines are involved in immune cell recruitment and have been associated with arthritic conditions whereby their levels correlate with inflammatory burden and concentrations of other cytokines that act on both immune cells and sensory neurons^54–57^. Therefore, it is plausible that BTB stimulates FLS to trigger release cytokine release that recruits immune cells and sensitises neurons.

### BTB exerts its pro-inflammatory effects via GPR65 expressing FLS

Results presented thus far, support the notion that BTB coordinates inflammatory joint pain in mice, most likely through its action on FLS resident in the knee joint which respond to the BTB by secreting inflammatory mediators that in turn sensitise DRG neurons. However, the requirement of GPR65 as the transducer of BTB remains to be proved. Given the lack of validated PS-GPCRs antagonists, GPR65 knockout (KO) mice^58^ were used to confirm the effects of BTB described with wild-type (WT) mice and cells. Firstly, FLS were cultured from KO mice; cells displayed no cAMP accumulation when challenged with BTB (Vehicle, 628.6 ± 129.3 pM, 10 µM BTB, 781.8 ± 808.8 pM, t = 0.739, df = 3, *p-adj =* 1.00; Fig. 5a) and an attenuated pH 6 response compared to that seen for WT FLS (Vehicle, 628.6 ± 129.3 pM, pH 6, 1.41 ± 0.12 nM, t = 12.385, *p-adj* = 0.007 ;pH 6 WT, 4.00 ± 0.16 nM, pH 6 KO, 1.41 ± 0.12 nM, t = -10.58, df = 3, *p-adj* = 0.002; Figs 4c, 5a). To determine whether GPR65 is necessary for the release of inflammatory mediators shown to sensitise DRG neurons, FLS cultured from WT or KO mice were stimulated with BTB overnight, conditioned media collected the following day was then incubated with naïve WT DRG neurons overnight before electrophysiological characterisation (Fig. 5b). The rheobase of DRG neurons cultured in the media from WT BTB-stimulated-FLS was markedly reduced compared to neurons cultured in the media from KO FLS (WT, 383.61 ± 44.55 pA, KO, 554.52 ± 44.98 pA, t = 2.670, df = 64.515, *p* = 0.0089; Fig. 5c-d). Whether DRG neurons were cultured in conditioned media from WT or KO FLS did not affect any other intrinsic or active property (Table 4). As before neurons incubated with the media from WT FLS stimulated with BTB exhibited larger voltage-gated inward current densities (WT, -539.00 ± 56.47 pA/pF, KO, -315.63 ± 49.97 pA/pF, t = 2.963, df = 19.708, *p* = 0.008; Fig. 5e-f), where minimal effect was seen on outward currents (WT, 547.18 ± 94.21 pA/pF, KO, 341.72 ± 63.78 pA/pF, t = -1.718, df = 17.575, *p* = 0.103; Fig. 5g-h).

**Figure 5.**
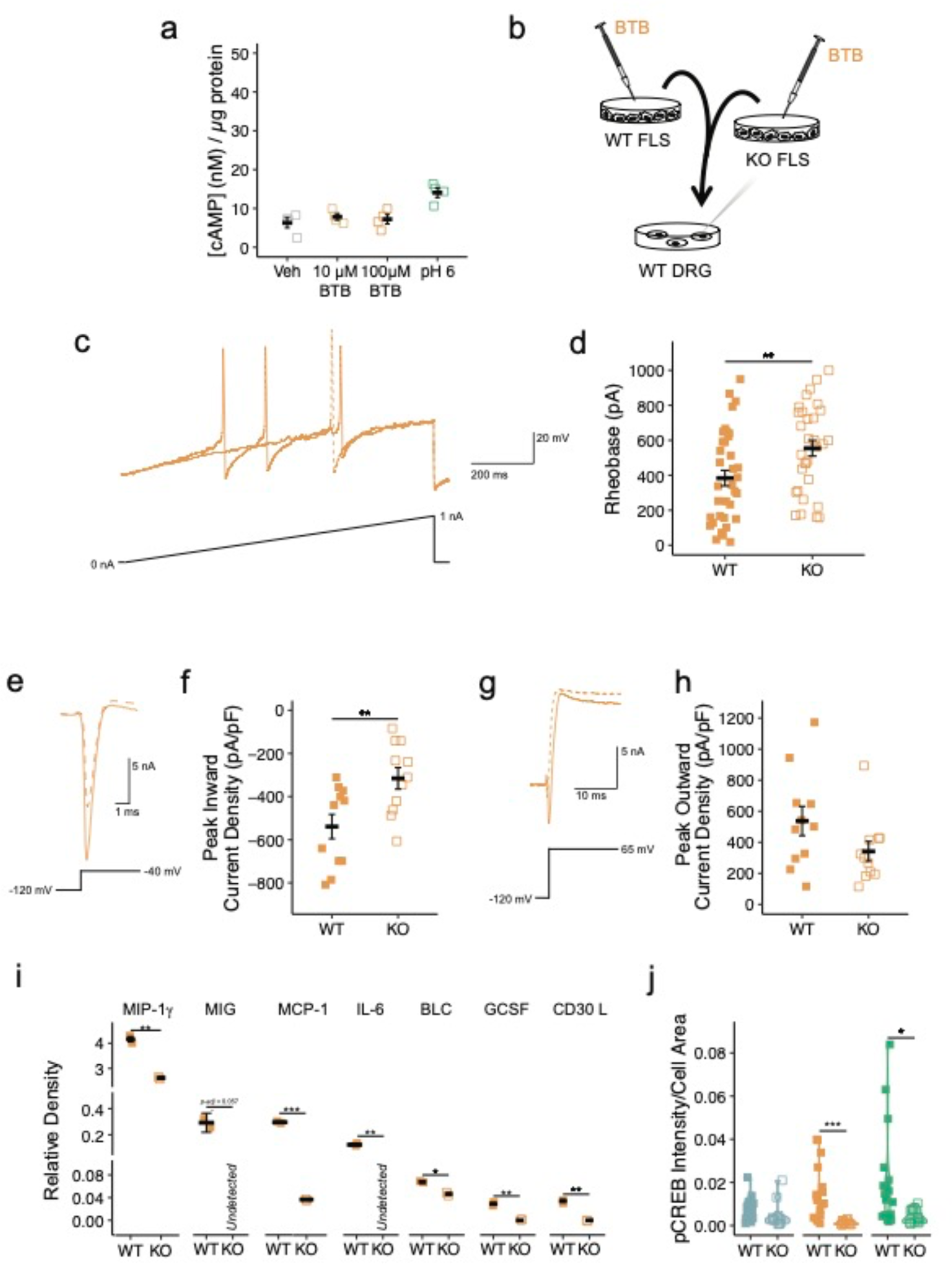
BTB exerts its pro-inflammatory effects via GPR65 expressing FLS. **(a)** Intracellular FLS cAMP concentration of FLS from GPR65 KO mice following stimulation with pH 7.4 vehicle, BTB or pH 6 solution. **(b)** WT (closed symbols/solid lines) and KO FLS (open symbols/dashed lines) were cultured overnight with BTB, the conditioned media collected 24-hours post-stimulation was then incubated with naïve DRG neurons overnight before electrophysiological characterisation. **(c)** Representative current clamp recordings of neurons of comparable capacitance, showing action potentials evoked by ramp injection of current (0– 1 nA, 1 s). **(d)** Stepwise current injections were used to determine the rheobase of sensory neurons. **(e)** Representative inward current trace from IV analyses, elicited by a -40 mV test pulse in cells of comparable capacitance. **(f)** Peak inward current density from IV analyses. **(g)** Representative outward current trace from IV analyses, elicited by a 65 mV test pulse in cells of comparable capacitance. **(h)** Peak outward current density from IV analyses. **(i)** Relative quantification of pro-inflammatory cytokines present in the conditioned media of WT and KO FLS following BTB stimulation. **(j)** phosphoCREB staining intensity of cultures of FLS from WT or KO animals following overnight stimulation with DMSO (blue), BTB (orange) or pH 6 (green). * *p / p-adj <* 0.05, ** *p / p-adj* < 0.01: **(d, f, h)** unpaired t-test. **(i-j)** Two-way ANOVA followed by Bonferroni-corrected post-hoc.

**Table 4:** Intrinsic and active properties of naïve dorsal root ganglion neurons cultured in conditioned media collected from WT or KO FLS stimulated with BTB overnight.

| **Current-clamp** | **WT (n = 36)** | **KO (n = 31)** |
| --- | --- | --- |
| Resting membrane potential (mV) | $-47.86 \pm 1.11$ | $-47.61 \pm 1.06$ |
| Capacitance (pF) | $51.91 \pm 6.73$ | $53.33 \pm 5.64$ |
| # APs at suprathreshold | $1.27 \pm 0.15$ (n = 15) | $2.25 \pm 0.78$ (n = 12) |
| Action potential amplitude (mV) | $82.06 \pm 4.69$ | $80.39 \pm 3.87$ |
| Half peak duration (ms) | $1.86 \pm 0.17$ | $2.06 \pm 0.20$ |
| Afterhyperpolarisation amplitude (mV) | $15.33 \pm 0.71$ | $16.24 \pm 0.68$ |
| Afterhyperpolarisation duration (ms) | $10.48 \pm 1.05$ | $10.91 \pm 1.29$ |
| **Voltage-clamp** | **WT (n = 11)** | **KO (n = 11)** |
| Inward: Erev | $67.77 \pm 9.55$ | $53.36 \pm 7.86$ |
| Inward: V1/2 | $-41.61 \pm 2.00$ | $-47.21 \pm 2.07$ |

Among the cytokines released in response to BTB-stimulation of WT FLS (Fig. 4k), reduced levels were found in the media of KO FLS following exposure to BTB (MIP-1γ: WT, 4.16 ± 0.10, KO, 2.62 ± 0.05, *p-adj* = 0.0058; MIG: WT, 0.29 ± 0.07, KO, undetected, *p-adj* = 0.0569; CD30 ligand: WT, 0.03 ± 0.002, KO, undetected, *p-adj* = 0.0037; MCP1: WT, 0.30 ± 0.01, KO, 0.04 ± 0.002, *p-adj* = 0.000713; Fig. 5i). The comparison of BTB-stimulation of WT vs KO FLS also revealed that BTB stimulation of GPR65 resulted in higher levels of other pro-inflammatory cytokines such as granulocyte colony-stimulating factor (GCSF; WT, 0.03 ± 0.002, KO, undetected, *p-adj* = 0.0041; Fig. 5i), interleukin 6 (IL-6; WT, 0.12 ± 0.01, KO, undetected, *p-adj* = 0.0094; Fig. 5i) and B lymphocyte chemoattractant (BLC; WT, 0.07 ± 0.002, KO, 0.05 ± 0.002, *p-adj* = 0.0156; Fig. 5i), among others; full analysis detailed in Table S2. GCSF, IL-6 and BLC all have established roles in immune cell recruitment and joint degeneration in arthritis^59–61^, IL-6 is additionally thought to directly sensitise sensory neurons to induce mechanical and thermal hypersensitivity^62,63^.These findings suggest that BTB activation of GPR65 on FLS might coordinate the expression and secretion of pro-inflammatory cytokines. To further explore the role of GPR65 in regulation of gene expression further, the ability of BTB to induce phosphorylation, and thus activation, of CREB, a transcription factor downstream of the Gα_s_/cAMP pathway was examined in FLS via immunostaining of phopsho-CREB (pCREB). Compared to KO FLS, higher pCREB intensity was observed when WT FLS were stimulated with BTB (WT, 0.015 ± 0.003, KO, 0.001 ± 0.000, *p-adj* = 0.00051; Fig. 5j). Importantly DMSO treatment had no effect on pCREB staining in FLS of either genotype (WT, 0.007 ± 0.001, KO, 0.005 ± 0.002, *p-adj* = 0.246; Fig. 5j). To begin to address whether GPR65 might influence the FLS transcriptome under more endogenous conditions, the effect of pH 6 treatment on FLS was also examined and resulted in higher pCREB intensity staining for WT vs KO FLS (WT, 0.020 ± 0.005, KO, 0.004 ± 0.001, *p-adj* = 0.0198; Fig. 5j). This suggests that the acidity of synovial fluid in arthritic conditions might modulate FLS via GPR65 to induce transcription machinery that results in the secretion of pro-inflammatory mediators and contribute to arthritic pathology.

### GPR65 KO mice do not develop joint inflammation or pain following intra-articular injection of BTB

To confirm that GPR65 is responsible for the pain-like behaviours observed following intra-articular BTB knee injection (Fig. 2), cohorts of WT and KO mice were injected with BTB into the knee joint, joint swelling and pain-like behaviours were then measured at baseline and 24-hours post-injection, coinciding with the peak of BTB-induced inflammation (Fig. 2b). Having previously studied females, studies were extended to include both sexes. As before, intra-articular BTB knee injection caused swelling in WT mice (knee width ratio: Baseline, 1.00 ± 0.01, 24-hours, 1.22 ± 0.02, t = -10.72, df = 13, *p-adj* < 0.0001; Fig. 6b), but no inflammatory response was evoked in KO mice (knee width ratio: Baseline, 1.00 ± 0.01, 24-hours, 1.01 ± 0.01, t = -0.982, df = 15, *p-adj* = 0.342; Fig. 6b). A difference in the extent of inflammation was seen for WT mice following injection of BTB based on sex, female mice experiencing greater inflammation (knee width ratio at 24-hours: Female, 1.26 ± 0.03, Male, 1.18 ± 0.01, t = 3.92, df = 6, *p-adj* = 0.008; Fig. 6b). The increased mechanical sensitivity of ipsilateral joints of WT mice (Baseline, 341.33 ± 13.16 g, 24-hours, 135.38 ± 13.87 g, t = 10.04, df = 13, *p-adj* < 0.0001; Fig. 6c) was not observed for KO mice (Baseline, 335.18 ± 11.18 g, 24-hours, 320.94 ± 10.81 g, t = 0.839, df = 15, *p-adj* = 0.415; Fig. 6c). Neither genotype showed any change in mechanical sensitivity of the contralateral joint (F(1,52) = 0.041, *p* = 0.84; Supplementary Fig. 4). Digging behaviours were also unaffected following BTB administration in KO mice (latency: Baseline, 37.13 ± 8.00 s, 24-hours, 52.44 ± 11.06 s, t = -1.09, df = 15, *p-adj =* 0.294; Fig 6d; duration: Baseline, 11.68 ± 1.99 s, 24-hours, 8.26 ± 1.26 s, t = 1.79, df = 15, *p-adj =* 0.094; Fig. 6e; burrows: Baseline, 4.06 ± 0.25, 24-hours, 4.00 ± 0.30, t = 0.169, df = 15, *p-adj* = 0.868; Fig. 6f). Deficits in digging were however seen 24-hours post-injection of WT mice (latency: Baseline, 24.00 ± 5.03 s, 24-hours, 112.43 ± 14.70 s, t = -7.55, df = 13, *p-adj* < 0.0001; Fig. 6d; duration: Baseline, 13.13 ± 2.14 s, 24-hours, 3.17 ± 0.91 s, t = 4.88, df = 13, *p-adj* = 0.0003; Fig. 6e; burrows: Baseline, 4.5 ± 0.25, 24-hours, 1.93 ± 0.36, t = 9.47, df = 13, *p-adj* < 0.0001; Fig. 6f). Despite the inflammatory response following injection of BTB to WT differing for females and males, no sex effect was seen for any behavioural readout (Mechanical threshold: F(1,52) = 2.84, *p-adj* = 0.098; Digging latency: F(1,52) = 0.132, *p-adj* = 0.718; Digging duration: F(1,52) = 0.183, *p-adj* = 0.67; No. Burrows: F(1,52) = 0.00, *p-adj* = 1.00). Data here support the hypothesis that GPR65 mediates sensory neuron hyperexcitability, joint inflammation and pain-like behaviours conferred by BTB administration and highlight the potential of GPR65 antagonists in the treatment of inflammatory conditions.

**Figure 6.**
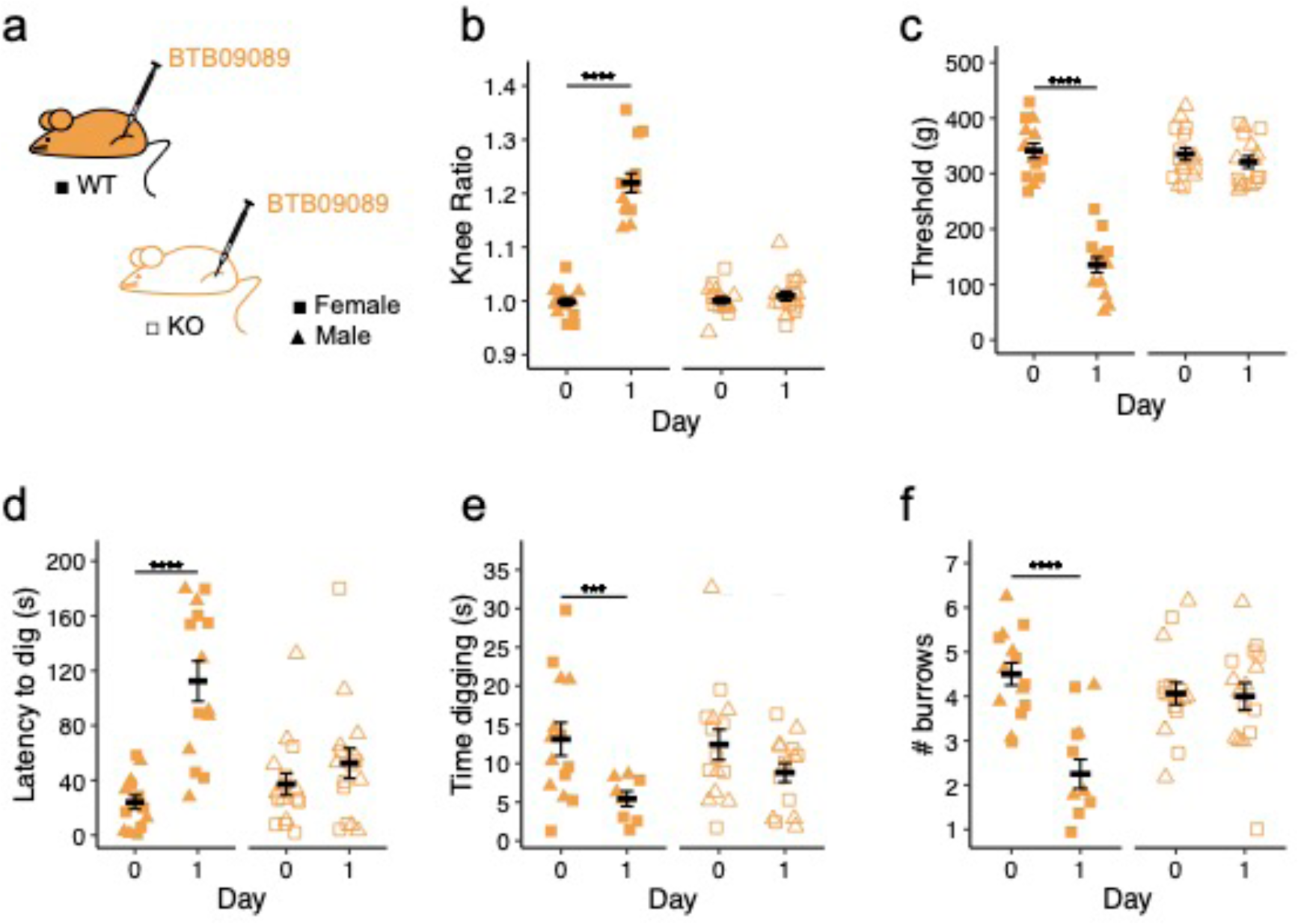
GPR65 KO mice do not develop joint inflammation or pain following intra-articular injection of BTB. **(a)** Wild-type (closed symbols) and GPR65 knockout (KO, open symbols) mice of either sex (females, square symbols; males, triangular symbols) received unilateral intra-articular injections of 100 µM BTB. **(b)** The ratio of the ipsilateral to contralateral knee width was calculated as a measure of the extent of inflammation. **(c)** Mechanical sensitivity of the injected knee joints was determined by pressure application measurement. The **(d)** latency to dig, **(e)** time spent digging and **(f)** number of burrows dug were also measured across experimental time. *** *p-adj* < 0.001, **** *p-adj* < 0.0001: repeated measures ANOVA followed by Bonferroni-corrected post-hoc. N = 14 wild-type mice (7 female, 7 male) and 16 KO mice (8 female, 8 male).

### Human FLS express GPR65 and arthritic synovial fluid samples activate GPR65

Having characterised a role of GPR65 in inflammatory joint pain in mice, the relevance of this signalling axis in human disease was explored next. Examining RNA sequencing data from Nanus and colleagues^53^, transcriptomic profiles of osteoarthritis (OA) patient synovium samples biopsied from sites reported as painful and non-painful revealed that GPR65 displayed one of the greatest increases in expression at painful sites, compared to non-painful sites of the proton-sensitive receptors for end-stage arthritic patients (Fold change: 3.83, *p-adj* = 0.117; Fig. 7a; Table S3). To further examine the potential role of GPR65 in the pathology of inflammatory arthritis in humans, FLS isolated from the painful sites of human OA patients were stimulated with BTB or DMSO for 24-hours, after which media were collected and later assayed for cytokine levels. Compared to FLS exposed to DMSO, higher levels of inflammatory mediators were detected in the media of human FLS stimulated with BTB. Most notable increases included IL-6 (BTB, 0.740 ± 0.014, DMSO, 0.054 ± 0.011, *p-adj* = 0.0068; Fig. 7b), already a therapeutic target in arthritis^64^ and IL-8 (BTB, 0.667 ± 0.001, DMSO, 0.066 ± 0.020, *p-adj* = 0.001; Fig. 7b), which is elevated in arthritic disease and has been shown to coordinate immune cell infiltration^65^. Human FLS stimulated with BTB released less MCP-1, compared to cells incubated with the vehicle, DMSO (BTB, 1.635 ± 0.033, DMSO, 2.610 ± 0.004, *p-adj* = 0.0012; Fig. 7b), which contrasts with findings from mouse FLS, where MCP1 was among the most upregulated cytokines (Fig. 4k). Possible explanations for this may include a species-related difference on the reliance of certain mediators in inflammatory programmes or an artefact of human FLS being isolated from OA patients, and so already influenced by the arthritic joint environment in addition to *in vitro* BTB stimulation, whereas FLS were obtained from healthy mice. Full analysis of the changes in cytokine levels following stimulation of human FLS are presented in Table S4.

**Figure 7.**
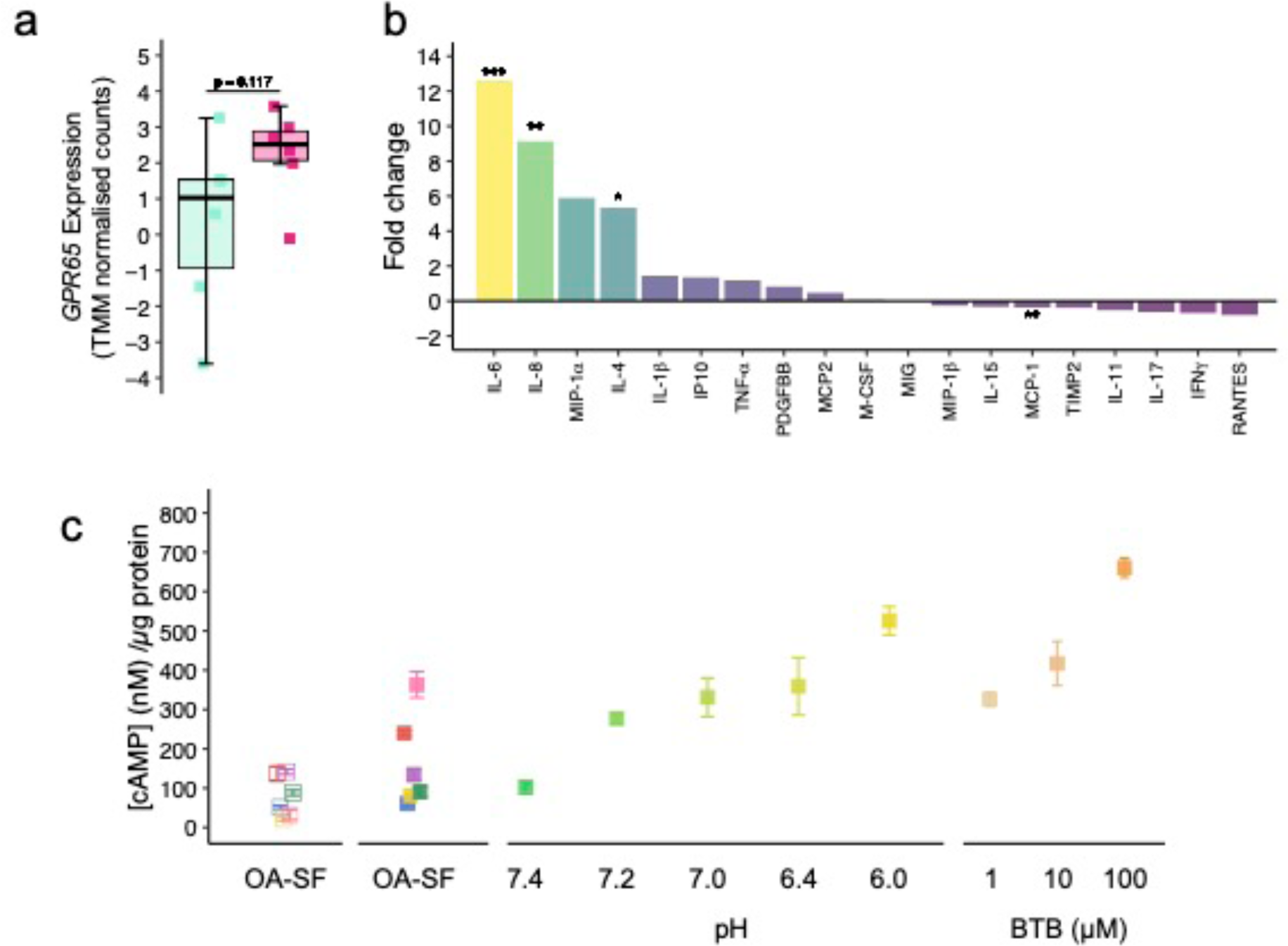
Human FLS express GPR65 and arthritic synovial fluid samples activate GPR65. **(a)** Expression of *GPR65* in synovium tissue from painful (pink) and non-painful (green) sites of end-stage OA patients, data from Nanus *et al*. (2021). **(b)** BTB-induced fold change in detection of inflammatory cytokines in conditioned media from stimulated human FLS. **(c)** Intracellular cAMP concentration of FlpIN CHO (open symbols) or mGPR65-CHO (closed symbols) cells, following stimulation with human OA synovial fluid samples, pH solutions or BTB. * *p-adj <* 0.05, ** *p-adj* < 0.01, *** *p-adj* < 0.001: **(a)** Published differentially expressed gene (DEG) data from Nanus *et al.* (2021) was filtered to identify DEGs using two-group statistical comparison for > 1.5 fold change and *p* < 0.05. **(b)** Two-way ANOVA followed by Bonferroni-corrected post-hoc.

Experiments presented thus far have relied on activation of GPR65 by BTB, which although selective among PS-GPCRs, is a synthetic tool, not an endogenous mediator. To address this, and further establish whether GPR65 contributes to inflammatory joint pain, samples of synovial fluid collected from OA patients (Table S5), were assessed for their ability to activate GPR65. The ability of samples to stimulate accumulation of cAMP in mGPR65-CHO cells was compared to responses seen in the parental Flp-IN CHO cell line (devoid of any endogenous proton-sensitive receptors). Of the six samples tested, three evoked a higher accumulation of cAMP in mGPR65-CHO cells than in parental cells, indicative of GPR65 activation (Fig. 7c). Alongside the synovial fluid samples, mGPR65-CHO cells were also challenged with acidic solutions or BTB, to compare the responses seen. Although the accumulation of cAMP observed cannot be equivocally pinned to engagement of GPR65 by protons in the synovial fluid, these data provide reassurance that the concentration of BTB selected for earlier studies was reasonable and still support a role of GPR65 in inflammatory arthritis pathology, suggesting further exploration as a therapeutic target is warranted.

## Discussion

Inflammatory conditions represent a significant burden on the quality of life for individuals affected, as well as having wider socioeconomic implications^3,66,67^. Accordingly, there is a pressing need to develop better and safer medications to remedy both the inflammation and pain associated with chronic diseases, due to the limitations of currently used medications. Here, a role of the PS-GPCR GPR65 in the development of inflammatory joint pain is presented, offering new insight into the cellular basis of inflammatory joint pain. The acidosis reported for inflammatory conditions, such as arthritis, makes a strong case for the involvement of proton-sensitive receptors in inflammation, however, delineating the roles of individual receptors is made difficult by the fact they are often co-expressed. To this end the selective GPR65 agonist BTB (Fig. 1) was used to explore its role in inflammatory joint pain. Following injection of BTB into the mouse knee joint an inflammatory response is initiated, which is evident from joint swelling. Mice also demonstrate behaviours consistent with pain, including mechanical hypersensitivity of the afflicted joint 24-hours after inflammation induction and longer lasting defects in digging behaviour reminiscent of spontaneous pain and inflammation-induced apathy (Fig. 2). The finding that stimulation of sensory neurons with BTB alone could not recreate the hyperexcitability seen of neurons that directly innervate BTB-injected joints, suggested that another cell type may be involved (Fig. 3). FLS resident in the knee joint express higher levels of GPR65 than any other PS-GPCR, and stimulation of FLS with BTB resulted in accumulation of intracellular cAMP (Fig. 4c), activation of the transcription factor CREB (Fig. 5h) and release of pro-inflammatory cytokines (Fig. 4k). Incubation of neurons in conditioned media taken from BTB-stimulated FLS lead to neuronal hyperexcitability and greater voltage-gated inward currents, akin to the effects seen for knee-innervating neurons isolated from mice injected with BTB (Figs. 3f-h, 4f-h). It is thus postulated that the inflammation and associated pain resulting from intra-articular BTB injection at least partially arises from activation of FLS resident in the joint, which are triggered to release mediators that can stimulate immune cells and sensory neurons in the vicinity to drive inflammation and pain.

Further evidence to support the specific involvement of GPR65 in inflammatory pain arise from the findings that GPR65 KO mice did not experience any inflammation or pain following injection of BTB (Fig. 6), FLS cultured from GPR65 KO mice did not exhibit BTB-induced cAMP accumulation (Fig. 5a) or release of mediators (Fig. 5j), and BTB-stimulated conditioned media from GPR65 KO FLS also failed to sensitise sensory neurons (Fig. 5d). These findings make use of the selective GPR65 agonist BTB, a convenient, but synthetic tool. The effect of protons, an endogenous GPR65 agonist, have also been explored throughout this study: BTB recapitulated most of the signalling effects of protons at GPR65 (Fig. 1) and protons were also shown to coordinate cAMP accumulation (Fig. 4c) and pCREB activation (Fig. 5k) for WT FLS, suggesting a similar mechanism may be initiated by natural acidosis which can occur during inflammatory arthritis. Furthermore, similar results were found for human FLS, and an ability of synovial fluid obtained from human OA patients to activate GPR65 was demonstrated. Integrating the findings reported here a potential mechanism for GPR65 in the mediation of arthritic pain may be postulated, whereby the acidosis of inflammation activates GPR65 expressed by FLS in the joint, which results in the production and secretion of inflammatory mediators which function to further recruit more immune cells, with mediators released by both FLS and the recruited immune cells capable of sensitising nearby sensory neurons to coordinate heightened nociception (Fig. 8).

**Figure 8.**
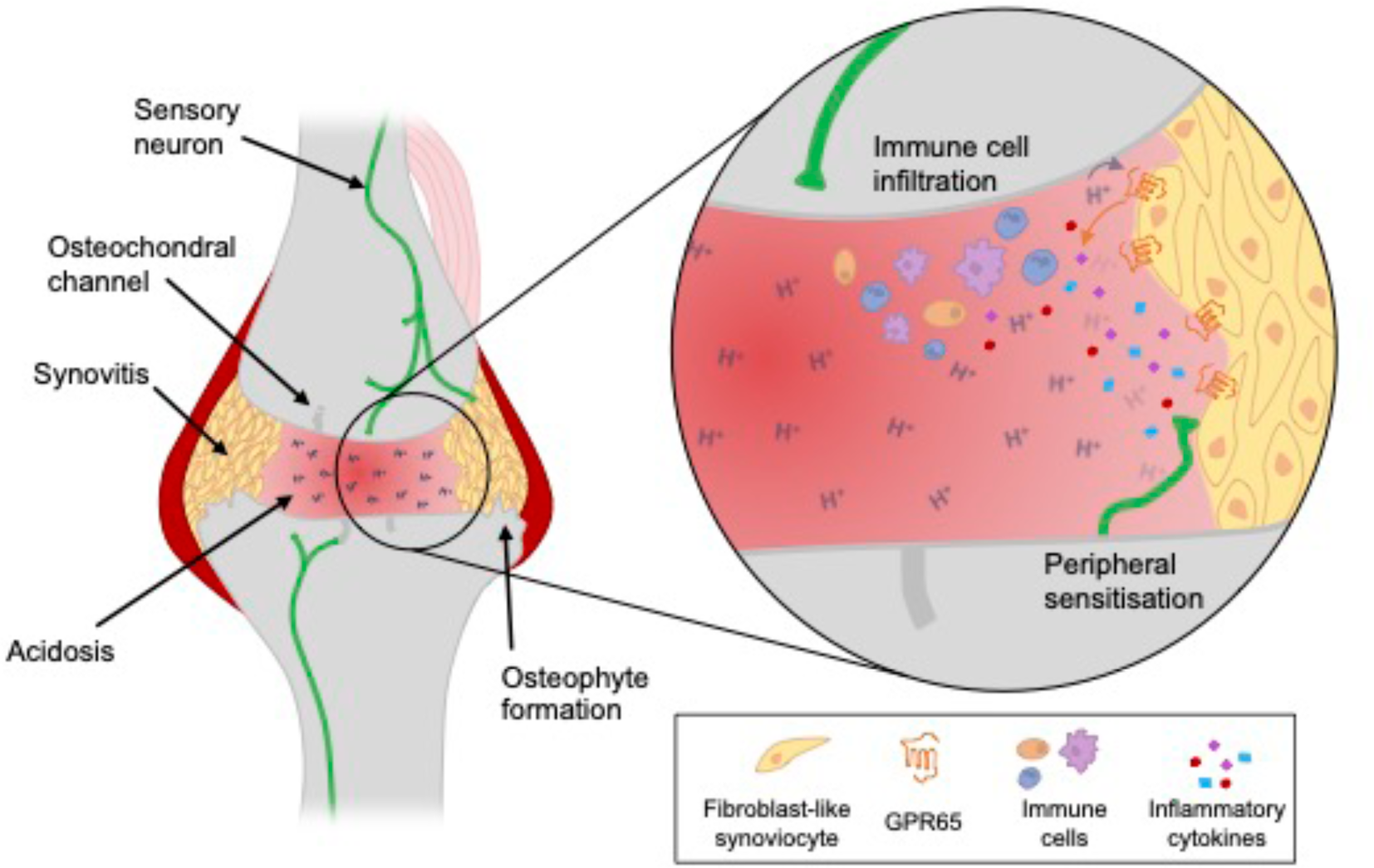
Activation of the proton-sensing GPCR, GPR65 on FLS contributes to inflammatory joint pain. Arthritic conditions are associated with localised acidosis of the joint environment and proliferation of FLS (synovitis). FLS express GPR65, are activated by the increased local concentration of protons (H^+^), leading to release of pro-inflammatory mediators capable of recruiting immune cells to the area, further driving inflammation. Mediators released by both FLS and immune cells act on sensory neurons that innervate the joint contributing to peripheral sensitisation and the increased pain associated with arthritis.

Stimulation of naïve cultures of sensory neurons with BTB alone was shown to be insufficient to recreate the hyperexcitability seen for neurons that project to BTB-injected joints (Fig. 3f). This might be explained by the very low expression of GPR65 by sensory neurons^68^ and thus the involvement of other cell types was explored. Data reported here suggesting a major role for FLS expressing GPR65 in establishing peripheral sensitisation of sensory neurons when these cells are studied in isolation (Figs. 4-5). Activation of GPR65 through injection of BTB into the knee joint also coordinated immune cell infiltration and an increase in pain-like behaviours (Figs. 2, 7). However, the KO mice used to attribute the inflammation and pain induced by BTB to GPR65 activation are global knockouts and it is thus impossible to resolve the contributions of individual cell types to the pathological changes observed. The involvement of other cell types cannot be overlooked, especially when immune cells, including T-cells and macrophages are also reported to express GPR65^23^. Given most of the cytokines found to increase in concentration following BTB-stimulation of FLS are principally associated with immune cell recruitment, the BTB-stimulated immune cell infiltration and secondary wave of mediator release that will likely ensue, and the dependency of this on GPR65, is deserving of further investigation. This might be addressed using more sophisticated tools for ablating GPR65 expression in specific cell types. Such an approach could add valuable cellular context to GPR65 activation, given there are reports of extracellular acidification reducing the ability of macrophages to produce TNF-α and IL-6 in a GPR65/PKA-dependent manner^69^.

The principle means by which BTB and GPR65 have been linked to inflammatory pain in the present study is through the release of inflammatory mediators. Additional disease context has been added from studies of FLS and synovial fluid samples obtained from human OA patients. However, OA is primarily considered a degenerative disorder, rather than outright inflammatory condition, which may offer some explanation to the apparent inability of three of the OA synovial fluid samples to engage GPR65, which may have been obtained from patients experiencing less inflammation at the time of collection. Unfortunately, synovial fluid volumes available in this study prevented measurement of their pH, but wide variation of the pH of synovial fluid samples obtained from patients with various forms of arthritis has been reported^38–41^. Accordingly, the next step should be to look at the involvement of GPR65 in conditions with a higher degree of inflammation, which might also enable an assessment of how the endogenous acidosis associated with such conditions contributes to pathology, a starting point could be the antigen-induced arthritis model, which has been shown to cause joint acidosis in rats^4^ and might offer a more translational assessment of the therapeutic potential of targeting GPR65.

Considering that chronic inflammatory conditions, would result in prolonged periods of acidosis and findings reported here which indicate that GPR65 internalises in response to acidic challenge (Fig. 1g), the contributions of GPR65 signalling from endosomal platforms to coordination of inflammatory pain is also worthy of further investigation, especially given the relevance of subcellular location on the ability of other GPCRs associated with inflammatory pain to coordinate nociception^70–72^. This phenomenon represents an extra layer of intrigue for GPR65, and indeed other PS-GCPRs, due to the progressive acidification of the endomembrane system, which would likely impact upon receptor activity. The work presented here in addition to these discussion points, suggest that development of GPR65 antagonists may hold therapeutic value in the treatment of inflammatory pain, the nature by which the receptor would be activated and contribute to disease in a pathological setting, i.e. localised acidosis, also means that delivery of any future agents active against the receptor may be enhanced through exploiting recent developments in pH-sensitive drug delivery^73^.

In summary, work described here has demonstrated that the PS-GPCR GRP65 is a regulator of inflammatory pain. The findings presented suggest that GPR65 coordinates inflammatory joint pain through FLS cells, which release inflammatory mediators in response to the GPR65 agonist BTB. These results provoke further questions regarding GPR65-driven inflammation and nociception, to allow better assessment of the receptor as a potential target for the development of novel anti-inflammatory and analgesic agents. Moreover, work here highlights the importance and potential value in further investigation of the wider PS-GPCR family in physiology and disease.

## Methods

*Please see Supplementary File 1 for comprehensive descriptions of methods*.

### Ethical approval

Ethical approval for obtaining human tissue and synovial fluid samples was granted by the UK National Research Ethics Committee (14/ES/1044 and REC 16/SS/0172). Consent was provided by all patients. All animal work was regulated in accordance with the United Kingdom Animal (Scientific Procedures) Act 1986 Amendment Regulations 2012 and was approved by the University of Cambridge Animal Welfare Ethical Review Body.

### Animals

Wildtype C57BL/6J mice (Envigo) and GPR65 KO mice^58^ were housed in groups of up to five per cage with enrichments and *ad libitum* access to food and water. The holding room was maintained at 21 °C and operated a 12-hour light/dark cycle. Mice were used at age 10-12 weeks.

### In vivo studies

Intra-articular injections, performed under anaesthesia, were made through the patella tendon. Fast Blue (Polysciences) was used to label knee-innervating neurons. BTB or DMSO was injected unilaterally to stimulate GPR65. Knee inflammation was measured using digital callipers. Mechanical sensitivity of the knee joint was assessed using pressure application measurement (Ugo Basile) and digging behaviour assessed as an ethological readout of pain and wellbeing^3^. Rotarod performance (Ugo Basile) was used to monitor mouse motor coordination. Behavioural analyses were carried out after blinding of experimental condition.

### Cell culture

Flp-In-CHO cells (Thermo Fisher) were used to generate stable PS-GPCR cell-lines. PS-GPCRs were cloned from mouse cDNA libraries using conventional restriction cloning methods. Expression of constructs by stable cells was confirmed through immunostaining and RT-PCR. Parental Flp-In-CHO cells were used as a control due to the lack of endogenous proton-sensitive receptor expression, or when experiments relied on transient transfection of tagged receptor constructs, i.e. bioluminescent resonance energy transfer (BRET) studies.

### Isolation and culture of mouse dorsal root ganglia (DRG) neurons

Following humane killing of mice, lumbar DRG (L2-L5) were isolated and incubated in collagenase (3mg, 15 min; Merck) before incubation in trypsin (3mg, 30 min; Merck). Mechanical trituration was used to dissociate neurons, which were then plated on poly-D-lysine and laminin glass-bottomed dishes (MaTek). Where DRG were taken from mice injected with BTB, neurons from the ipsilateral and contralateral sides were collected and cultured separately. Overnight incubation of DRG neurons with BTB or DMSO was to a total volume of 2 ml, Stimulation of naïve neurons with BTB or DMSO was to a total volume of 2 ml, and 300 µl for FLS conditioned media.

### Isolation and culture of mouse FLS

As conducted previously^52^, mice patellae were exposed by resection of the quadriceps muscle and briefly washed in PBS before transfer to culture media. After a week of outgrowth patellae were discarded and cells passaged, FLS were enriched through three subsequent passages before experimentation.

### Isolation and culture of humanFLS

FLS were isolated from joint synovial tissues collected peri-operatively from consenting OA patients following elective total joint replacement. Synovial tissue was prepared as previously described^74^ in complete fibroblast media, and isolated FLS were enriched and maintained till 70% confluent up to passage 4 before plating for experiments.

### Cell line signalling assays

cAMP accumulation and ERK phosphorylation were quantified using LANCE Ultra TR-FRET reagents (Perkin Elmer). The Ca^2+^-sensitive dye, Fluo-4 (Thermo Fisher) was used to study Ca^2+^ mobilisation. β-arrestin recruitment and receptor internalisation were assessed by bystander BRET assays.

*cAMP accumulation in FLS* FLS were stimulated for 15 minutes followed by cell lysis. To account for the variability in cell number in experiments, cAMP concentration in lysates (inferred from a cAMP standard curve run in parallel, using LANCE Ultra TR-FRET reagents) were normalised to the total protein concentration in each lysate (assessed by Bradford assay).

*qPCR* RNA was isolated from FLS using TRIzol (Merck) and an RNA clean up and concentrator kit (Zymo Research). High-capacity reverse transcription reagents (Applied Biosystems) were used to synthesis cDNA and gene expression was assayed using TaqMan probes (Thermo Fisher). Relative expression was calculated as 2^-ýCt^, where ΔCt is the difference in the Ct value obtained for the gene of interest minus that of the housekeeping gene, *18SrRNA*.

### Electrophysiology

Step-wise depolarisation (Δ10 pA, 50 ms) was used to determine the rheobase. The activity of macroscopic voltage-gated channels was also assessed in voltage-clamp mode with appropriate series resistance compensation. Neurons were held at -120 mV for 150 ms before stepping to the test potential (−60 mV – 55 mV in 5 mV increments) for 40 ms and returning to a holding potential of -60 mV for 200 ms between sweeps. Peak inward and outward currents were normalised to cell capacitance. Fast Blue, knee-innervating, neurons were identified by LED excitation at 365 nm (Cairn Research).

### Cytokine analyses

FLS were stimulated with 100 µM BTB or 0.1% (v/v) DMSO in serum-depleted media for 24-hours after which conditioned media were collected. Cytokine levels in collected media were assayed with Mouse or Human Inflammatory Antibody Arrays (Abcam) as appropriate.

### Immunostaining

FLS were fixed by exposure to ice-cold 100% methanol and stained for CDH-11 and phospho-CREB. The nucleus was stained with DAPI. Cells were imaged with a Leica SP5 laser-scanning confocal microscope and 63x oil objective. The intensity of pCREB staining was analysed in ImageJ.

### Data analysis and statistics

Data are presented as mean ± standard error of the mean, the number of biological and technical replicates are detailed in individual figure legends. Appropriate analyses were selected according to the number of factors being compared and whether data met the assumptions for parametric analyses, the statistical tests employed are stated in corresponding figure legends. All analyses were performed in R, and *p* values < 0.05 considered significant.

## Acknowledgements

LAP was supported by the University of Cambridge BBSRC Doctoral Training Programme (BB/MO11194/1) and Corpus Christi College, University of Cambridge. LAP and ESS acknowledge funding from the MRC (MR/W002426/1). RHR was supported by an AstraZeneca PhD Studentship (G104108). HH was supported by a BBSRC/GSK iCASE PhD Studentship (BB/V509528/1). SNW and SWJ acknowledge funding from the MRC (MR/W026961/1) and Versus Arthritis (21530). GL was supported by a Royal Society Industry Fellowship (NF\R2\212001).

## Author contributions

Conception and design: LAP and ESS. Preparation and provision of human samples, mouse lines and reagents: LAP, SNW, GL, LY and SWJ. Acquisition and analysis of data: LAP, RHR, HH. Interpretation of data: LAP and ESS. Writing of manuscript: LAP and ESS. All authors approved the final version of this manuscript.

## Data availability

Data sets supporting the conclusions of this article are available in University of Cambridge Apollo Repository (https://doi.org/10.17863/CAM.106374).

## References

1. Ghouri, A. & Conaghan, P. G. Treating osteoarthritis pain: recent approaches using pharmacological therapies. Clin. Exp. Rheumatol. 37 Suppl 120, 124–129 (2019).

2. Radu, A.-F. & Bungau, S. G. Management of Rheumatoid Arthritis: An Overview. Cells 10, 2857 (2021).

3. Pattison, L. A. et al. Digging deeper into pain: an ethological behavior assay correlating well-being in mice with human pain experience. Pain (2023) doi:10.1097/j.pain.0000000000003190.

4. Andersson, S. E., Lexmüller, K., Johansson, A. & Ekström, G. M. Tissue and intracellular pH in normal periarticular soft tissue and during different phases of antigen induced arthritis in the rat. J. Rheumatol. 26, 2018–24 (1999).

5. Sasaki, Y., Hada, R., Nakajima, H., Fukuda, S. & Munakata, A. Improved localizing method of radiopill in measurement of entire gastrointestinal pH profiles: colonic luminal pH in normal subjects and patients with Crohn’s disease. Am. J. Gastroenterol. 92, 114–8 (1997).

6. Menkin, V. & Warner, C. R. Studies on Inflammation: XIII. Carbohydrate Metabolism, Local Acidosis, and the Cytological Picture in Inflammation. Am. J. Pathol. 13, 25–44.1 (1937).

7. Woo, Y. C., Park, S. S., Subieta, A. R. & Brennan, T. J. Changes in Tissue pH and Temperature after Incision Indicate Acidosis May Contribute to Postoperative Pain. Anesthesiology 101, 468–475 (2004).

8. Zwieten, R. van, Wever, R., Hamers, M. N., Weening, R. S. & Roos, D. Extracellular proton release by stimulated neutrophils. J. Clin. Investig. 68, 310–313 (1981).

9. Riemann, A. et al. Acidosis differently modulates the inflammatory program in monocytes and macrophages. Biochim. Biophys. Acta (BBA) - Mol. Basis Dis. 1862, 72–81 (2016).

10. Gregory, N. S., Whitley, P. E. & Sluka, K. A. Effect of Intramuscular Protons, Lactate, and ATP on Muscle Hyperalgesia in Rats. PLoS ONE 10, e0138576 (2015).

11. Steen, K. H., Steen, A. E. & Reeh, P. W. A dominant role of acid pH in inflammatory excitation and sensitization of nociceptors in rat skin, in vitro. J. Neurosci. : Off. J. Soc. Neurosci. 15, 3982–9 (1995).

12. Jones, N. G., Slater, R., Cadiou, H., McNaughton, P. & McMahon, S. B. Acid-Induced Pain and Its Modulation in Humans. J. Neurosci. 24, 10974–10979 (2004).

13. Ugawa, S. et al. Amiloride-blockable acid-sensing ion channels are leading acid sensors expressed in human nociceptors. J. Clin. Investig. 110, 1185–1190 (2002).

14. Pattison, L. A., Callejo, G. & Smith, E. S. J. Evolution of acid nociception: ion channels and receptors for detecting acid. Philos. Trans. R. Soc. B: Biol. Sci. 374, 20190291 (2019).

15. Murata, N. et al. Inhibition of superoxide anion production by extracellular acidification in neutrophils. Cell. Immunol. 259, 21–26 (2009).

16. Riemann, A. et al. Acidic environment activates inflammatory programs in fibroblasts via a cAMP–MAPK pathway. Biochim. Biophys. Acta (BBA) - Mol. Cell Res. 1853, 299–307 (2015).

17. Huang, C.-W. et al. Nociceptors of dorsal root ganglion express proton-sensing G-protein-coupled receptors. Mol. Cell. Neurosci. 36, 195–210 (2007).

18. Chen, Y.-J., Huang, C.-W., Lin, C.-S., Chang, W.-H. & Sun, W.-H. Expression and Function of Proton-Sensing G-Protein-Coupled Receptors in Inflammatory Pain. Mol. Pain 5, 1744-8069-5–39 (2009).

19. Kottyan, L. C. et al. Eosinophil viability is increased by acidic pH in a cAMP- and GPR65-dependent manner. Blood 114, 2774–2782 (2009).

20. Kringel, D., Kaunisto, M. A., Kalso, E. & Lötsch, J. Machine-learned analysis of the association of next-generation sequencing-based genotypes with persistent pain after breast cancer surgery. Pain 160, 2263–2277 (2019).

21. Wang, Y. et al. The Proton-activated Receptor GPR4 Modulates Intestinal Inflammation. J. Crohn’s Colitis 12, 355–368 (2018).

22. Vallière, C. de et al. G Protein-coupled pH-sensing Receptor OGR1 Is a Regulator of Intestinal Inflammation. Inflamm. Bowel Dis. 21, 1269–1281 (2015).

23. Tcymbarevich, I. V. et al. The impact of the rs8005161 polymorphism on G protein-coupled receptor GPR65 (TDAG8) pH-associated activation in intestinal inflammation. BMC Gastroenterol. 19, 2 (2019).

24. Hu, W.-P., Zeng, Y.-Y., Zuo, Y.-H. & Zhang, J. Identification of novel candidate genes involved in the progression of emphysema by bioinformatic methods. Int. J. Chronic Obstr. Pulm. Dis. 13, 3733–3747 (2018).

25. Huang, W.-Y., Dai, S.-P., Chang, Y.-C. & Sun, W.-H. Acidosis Mediates the Switching of Gs-PKA and Gi-PKCε Dependence in Prolonged Hyperalgesia Induced by Inflammation. PLoS ONE 10, e0125022 (2015).

26. Lin, S.-H. et al. Involvement of TRPV1 and TDAG8 in Pruriception Associated with Noxious Acidosis. J. Investig. Dermatol. 137, 170–178 (2017).

27. Hsieh, W.-S., Kung, C.-C., Huang, S.-L., Lin, S.-C. & Sun, W.-H. TDAG8, TRPV1, and ASIC3 involved in establishing hyperalgesic priming in experimental rheumatoid arthritis. Sci. Rep. 7, 8870 (2017).

28. Dai, S.-P. et al. TDAG8 deficiency reduces satellite glial number and pro-inflammatory macrophage number to relieve rheumatoid arthritis disease severity and chronic pain. J. Neuroinflammation 17, 170 (2020).

29. Dai, S.-P. et al. TDAG8 involved in initiating inflammatory hyperalgesia and establishing hyperalgesic priming in mice. Sci. Rep. 7, 41415 (2017).

30. Lin, R. et al. GPR65 promotes intestinal mucosal Th1 and Th17 cell differentiation and gut inflammation through downregulating NUAK2. Clin. Transl. Med. 12, e771 (2022).

31. Zhang, K. et al. Targeting GPR65 alleviates hepatic inflammation and fibrosis by suppressing the JNK and NF-κB pathways. Mil. Méd. Res. 10, 56 (2023).

32. Hardin, M. et al. The clinical and genetic features of COPD-asthma overlap syndrome. Eur. Respir. J. 44, 341–350 (2014).

33. Yuan, Y. et al. Genetic polymorphisms of G protein-coupled receptor 65 gene are associated with ankylosing spondylitis in a Chinese Han population: A case-control study. Hum. Immunol. 80, 146–150 (2019).

34. Xie, L. et al. pH and Proton Sensor GPR65 Determine Susceptibility to Atopic Dermatitis. J. Immunol. 207, 101–109 (2021).

35. Wang, J.-Q. et al. TDAG8 is a proton-sensing and psychosine-sensitive G-protein-coupled receptor. J. Biol. Chem. 279, 45626–33 (2004).

36. Im, D.-S., Heise, C. E., Nguyen, T., O’Dowd, B. F. & Lynch, K. R. Identification of a Molecular Target of Psychosine and Its Role in Globoid Cell Formation. J. Cell Biol. 153, 429– 434 (2001).

37. Onozawa, Y. et al. Activation of T cell death-associated gene 8 regulates the cytokine production of T cells and macrophages in vitro. Eur. J. Pharmacol. 683, 325–331 (2012).

38. Goldie, I. & Nachemson, A. Synovial pH in Rheumatoid Knee-Joints I. The Effect of Synovectomy. Acta Orthop. Scand. 40, 634–641 (2009).

39. Ward, T. T. & Steigbigel, R. T. Acidosis of synovial fluid correlates with synovial fluid leukocytosis. Am. J. Med. 64, 933–936 (1978).

40. Treuhaft, P. S. & McCarty, D. J. Synovial fluid pH, lactate, oxygen and carbon dioxide partial pressure in various joint diseases. Arthritis Rheum. 14, 475–484 (1971).

41. Roman, M. D. et al. Assesment of Synovial Fluid pH in Osteoarthritis of the HIP and Knee. Rev. Chim. 68, 1242–1244 (2017).

42. Safiri, S. et al. Global, regional and national burden of osteoarthritis 1990-2017: a systematic analysis of the Global Burden of Disease Study 2017. Ann. Rheum. Dis. 79, 819– 828 (2020).

43. Safiri, S. et al. Global, regional and national burden of rheumatoid arthritis 1990–2017: a systematic analysis of the Global Burden of Disease study 2017. Ann. Rheum. Dis. 78, 1463 (2019).

44. Smith, E. St. J. Advances in understanding nociception and neuropathic pain. J. Neurol. 265, 231–238 (2018).

45. Pinho-Ribeiro, F. A., Verri, W. A. & Chiu, I. M. Nociceptor Sensory Neuron–Immune Interactions in Pain and Inflammation. Trends Immunol. 38, 5–19 (2017).

46. Li, Z., Huang, Z. & Bai, L. Cell Interplay in Osteoarthritis. Front. Cell Dev. Biol. 9, 720477 (2021).

47. Pattison, L. A., Krock, E., Svensson, C. I. & Smith, E. S. J. Cell–cell interactions in joint pain: rheumatoid arthritis and osteoarthritis. Pain 162, 714–717 (2021).

48. Nandakumar, K. S., Fang, Q., Ågren, I. W. & Bejmo, Z. F. Aberrant Activation of Immune and Non-Immune Cells Contributes to Joint Inflammation and Bone Degradation in Rheumatoid Arthritis. Int. J. Mol. Sci. 24, 15883 (2023).

49. Han, D. et al. The emerging role of fibroblast-like synoviocytes-mediated synovitis in osteoarthritis: An update. J. Cell. Mol. Med. 24, 9518–9532 (2020).

50. Tsaltskan, V. & Firestein, G. S. Targeting fibroblast-like synoviocytes in rheumatoid arthritis. Curr. Opin. Pharmacol. 67, 102304 (2022).

51. Bai, Z. et al. Synovial fibroblast gene expression is associated with sensory nerve growth and pain in rheumatoid arthritis. Sci. Transl. Med. 16, eadk3506 (2024).

52. Chakrabarti, S. et al. Sensitization of knee-innervating sensory neurons by tumor necrosis factor-α-activated fibroblast-like synoviocytes: an in vitro, coculture model of inflammatory pain. Pain 161, 2129–2141 (2020).

53. Nanus, D. E. et al. Synovial tissue from sites of joint pain in knee osteoarthritis patients exhibits a differential phenotype with distinct fibroblast subsets. EBioMedicine 72, 103618 (2021).

54. Lean, J. M., Murphy, C., Fuller, K. & Chambers, T. J. CCL9/MIP-1gamma and its receptor CCR1 are the major chemokine ligand/receptor species expressed by osteoclasts. J. Cell. Biochem. 87, 386–93 (2002).

55. Tsubaki, T. et al. Accumulation of plasma cells expressing CXCR3 in the synovial sublining regions of early rheumatoid arthritis in association with production of Mig/CXCL9 by synovial fibroblasts. Clin. Exp. Immunol. 141, 363–371 (2005).

56. Barbieri, A. et al. Characterization of CD30/CD30L+ Cells in Peripheral Blood and Synovial Fluid of Patients with Rheumatoid Arthritis. J. Immunol. Res. 2015, 729654 (2015).

57. Harigai, M. et al. Monocyte Chemoattractant Protein-1 (MCP-1) in Inflammatory Joint Diseases and Its Involvement in the Cytokine Network of Rheumatoid Synovium. Clin. Immunol. Immunopathol. 69, 83–91 (1993).

58. Radu, C. G. et al. Normal Immune Development and Glucocorticoid-Induced Thymocyte Apoptosis in Mice Deficient for the T-Cell Death-Associated Gene 8 Receptor. Mol. Cell. Biol. 26, 668–677 (2006).

59. Lawlor, K. E. et al. Critical role for granulocyte colony-stimulating factor in inflammatory arthritis. Proc. Natl. Acad. Sci. 101, 11398–11403 (2004).

60. Fujimoto, M. et al. Interleukin-6 blockade suppresses autoimmune arthritis in mice by the inhibition of inflammatory Th17 responses. Arthritis Rheum. 58, 3710–3719 (2008).

61. Bugatti, S. et al. High expression levels of the B cell chemoattractant CXCL13 in rheumatoid synovium are a marker of severe disease. Rheumatology 53, 1886–1895 (2014).

62. Brenn, D., Richter, F. & Schaible, H. Sensitization of unmyelinated sensory fibers of the joint nerve to mechanical stimuli by interleukin-6 in the rat: An inflammatory mechanism of joint pain. Arthritis Rheum. 56, 351–359 (2007).

63. Obreja, O., Schmelz, M., Poole, S. & Kress, M. Interleukin-6 in combination with its soluble IL-6 receptor sensitises rat skin nociceptors to heat, in vivo. Pain 96, 57–62 (2002).

64. Burmester, G. R. et al. Tocilizumab in early progressive rheumatoid arthritis: FUNCTION, a randomised controlled trial. Ann. Rheum. Dis. 75, 1081 (2016).

65. Endo, H., Akahoshi, T., Takagishi, K., Kashiwazaki, S. & Matsushima, K. Elevation of interleukin-8 (IL-8) levels in joint fluids of patients with rheumatoid arthritis and the induction by IL-8 of leukocyte infiltration and synovitis in rabbit joints. Lymphokine cytokine Res. 10, 245–52 (1991).

66. Breivik, H., Collett, B., Ventafridda, V., Cohen, R. & Gallacher, D. Survey of chronic pain in Europe: Prevalence, impact on daily life, and treatment. Eur. J. Pain 10, 287–287 (2006).

67. Jacobs, P., Bissonnette, R. & Guenther, L. C. Socioeconomic burden of immune-mediated inflammatory diseases--focusing on work productivity and disability. J. Rheumatol. Suppl. 88, 55–61 (2011).

68. Zeisel, A. et al. Molecular Architecture of the Mouse Nervous System. Cell 174, 999–1014.e22 (2018).

69. Mogi, C. et al. Involvement of Proton-Sensing TDAG8 in Extracellular Acidification-Induced Inhibition of Proinflammatory Cytokine Production in Peritoneal Macrophages. J. Immunol. 182, 3243–3251 (2009).

70. Jensen, D. D. et al. Neurokinin 1 receptor signaling in endosomes mediates sustained nociception and is a viable therapeutic target for prolonged pain relief. Sci. Transl. Med. 9, (2017).

71. Yarwood, R. E. et al. Endosomal signaling of the receptor for calcitonin gene-related peptide mediates pain transmission. Proc. Natl. Acad. Sci. 114, 12309–12314 (2017).

72. Jimenez-Vargas, N. N., et al. Protease-activated receptor-2 in endosomes signals persistent pain of irritable bowel syndrome. Proc. Natl. Acad. Sci. 115, E7438–E7447 (2018).

73. Ramírez-García, P. D. et al. A pH-responsive nanoparticle targets the neurokinin 1 receptor in endosomes to prevent chronic pain. Nat. Nanotechnol. 14, 1150–1159 (2019).

74. Wijesinghe, S. N. et al. Obesity defined molecular endotypes in the synovium of patients with osteoarthritis provides a rationale for therapeutic targeting of fibroblast subsets. Clin. Transl. Med. 13, e1232 (2023).

