## Supplemental Information for "Activation of the proton-sensing GPCR, GPR65 on fibroblast-like synoviocytes contributes to inflammatory joint pain"

### Materials and Methods

*Ethical approval* Ethical approval for obtaining human tissue and synovial fluid samples was granted by the UK National Research Ethics Committee (14/ES/1044 and REC 16/SS/0172). Consent was provided by all patients. All animal work was regulated in accordance with the United Kingdom Animal (Scientific Procedures) Act 1986 Amendment Regulations 2012 and was approved by the University of Cambridge Animal Welfare Ethical Review Body.

*Agonists and activators* The GPR65 agonist, BTB09089 (BTB) was synthesised by Otava Chemicals and reconstituted in DMSO (Sigma Aldrich). Stocks of psychosine, a glycosphingolipid also reported to activate GPR65 (Sigma Aldrich) were prepared in chloroform:methanol (1:1). Forskolin (FSK, Sigma Aldrich), an adenylyl cyclase activator, was prepared in DMSO. The phorbol ester, phorbol 12,13-dibutyrate (PDBu, Tocris), an activator of mitogen-activated kinases, was prepared in DMSO. Ionomycin, a $Ca^{2+}$ ionophore (Cayman Chemicals) were prepared in ethanol. Coelenterazine H (Nanolight Technologies), a substrate of the Renilla luciferase, were prepared in 100% methanol.

*Animals* C57BL/6J wild-type (WT, Envigo) and GPR65 KO mice[1] were housed in groups of up to five per cage with enrichments and *ad libitum* access to food and water. The holding room was maintained at 21 °C and operated a 12-hour light/dark cycle. Mice were used at age 10-12 weeks. Throughout animal studies daily monitoring was performed: general welfare (signs of inappetence, coat condition and mobility) was observed and animal weight and joint inflammation (measured with digital Vernier callipers) were measured to ensure that the humane endpoints (excessive weight loss or joint swelling).

*Intra-articular injections* Intra-articular injections of mouse knee joints were performed under anaesthesia (ketamine, 100 mg/kg and xylazine, 10 mg/kg administered intra-peritoneally). After complete loss of consciousness injections were made through the patella tendon using a Hamilton syringe and 30 g needle. To label knee-innervating sensory neurons, 1.5 µl of the retrograde tracer Fast Blue (FB, 2% in saline, Polysciences) was injected. To activate GPR65 10 µl 100 µM BTB was injected unilaterally (ipsilateral knee randomly determined). Control cohorts were injected with 10 µl 0.1% (v/v) DMSO (Sigma Aldrich).

*Behavioural experiments* Before assessing any behaviours animals were acclimatised in behavioural testing rooms for at least 30-minutes each day of testing. The room used, time of testing (2-4 hours into the start of the light cycle of animals' holding room), and experimenters present were kept constant for each individual study to reduce confounds on observed behaviour.

*Behaviour – Digging* After acclimatization to the behaviour room and experimenters, mice were individually transferred to testing cages (49 × 10 × 12 cm) set up with ~4 cm of tightly packed fine-grain aspen midi 8/20 wood chip bedding (LBS Biotechnology), and up to 5 cages were tested simultaneously. Mice were allowed 3 minutes to explore testing cages without interference, under video surveillance. Training sessions were carried out the day before baseline behaviours were assessed to allow mice to gain familiarity with the setup. Training consisted of mice being placed in test cages as per a standard test. On the rare occasion that an animal did not dig, the individual was placed into a test cage with another animal (from the same home cage) that did meet this criterion, and a second training dig was conducted after a 30-minute break. Separate cages were used to test female and male mice, although when both sexes were studied, both were present in the behaviour room for the duration of testing. After testing sessions, the number of visible burrows remaining at the end of the 3-minute test was recorded. Digging behaviour was further analysed offline via video playback at the conclusion of each study, after blinding of all videos. The latency of mice to begin digging was noted and the total time mice spent digging was scored independently by at least 2 observers; because scores were well correlated ($R^2$ = 0.937), an average is reported as the time spent digging.

*Behaviour – RotaRod* A RotaRod apparatus (Ugo Basile 7650) was used to assess the motor coordination of mice. Animals were allowed 1 minute to acclimatize to the apparatus on a slow setting (7 revolutions per minute, rpm) before an accelerating program (7-40 rpm over 5 minutes) was started. Mice were first placed on the RotaRod after being trained in the digging assay to gain some familiarity with the apparatus. Tests were video-recorded, and mice were removed from the RotaRod if they fell, after 2 consecutive passive rotations or after 6 minutes of the accelerating program, whichever occurred first. Analysis of videos was performed at the conclusion of studies, and after blinding, the latency to passive rotation or fall was scored by a single experimenter.

*Behaviour – Pressure Application Measurement* As the only evoked pain behaviour tested, mechanical threshold was always assessed last, so as not to let recent infliction of a painful stimuli impact on the spontaneous behavioural assessments. Mechanical sensitivity of the knee joint was assessed using a pressure application measurement device (Ugo Basile). Animals were scruffed by 1 experimenter and presented to a second experimenter, blind to the animal's identity. While lightly holding the hind paw, to maintain a slight bend in the knee of the animal, the second experimenter applied gradual force to each knee joint, squeezing each joint medially with the force transducer. The withdrawal threshold was recorded when an animal withdrew the limb being tested or after 450g force (the upper limit threshold) was applied. Each animal was tested twice per time point, with a short break between tests; withdrawal force is reported as an average of the 2 measurements taken at each time point. Pressure application measurements were taken by a single experimenter for mice injected with BTB to be used for electrophysiology experiments 24-hours post-injection, thus the scuffing and application of force was slightly different for these experiments, which were also unable to be blinded.

*Isolation and culture of mouse dorsal root ganglia (DRG)* Mice were sacrificed by cervical dislocation before the lumbar DRG (L2 – L5) were collected in dissociation medium (L-15 + GlutaMAX growth media supplemented with 24 mM $NaHCO_3$; Life Technologies). For experiments on cells from mice which underwent unilateral injection of an inflammatory substance (carrageenan or BTB), DRG from the injected (ipsilateral, ipsi.) and non-injected (contralateral, contra.) sides were collected separately and underwent the rest of the culturing process independently. When naïve mice were used, lumbar DRG from both sides of each mouse were pooled and cultured together. Once dissected, DRG were incubated in dissociation media containing 1 mg/ml type 1A collagenase (Sigma-Aldrich) and 6 mg/ml BSA for 15 min at 37 °C, 5% $CO_2$, before a further 30 min incubation in dissociation media containing 1 mg/ml trypsin (Sigma-Aldrich) and 6 mg/ml BSA. DRG were then suspended in culture media (L-15 + GlutaMAX growth media supplemented with 10% (v/v) foetal bovine serum (FBS, Sigma Aldrich), 24 mM $NaHCO_3$ 38 mM glucose and 2% (v/v) penicillin/streptomycin (Life Technologies)) before several rounds of mechanical trituration and brief centrifugation (160g, 30 s). After sufficient trituration, dissociated cells were pelleted (160g, 5 min), resuspended in culture media and plated on poly-D-lysine/laminin coated glass coverslips (BD Biosciences) or glass-bottomed dishes (MatTek) and incubated at 37 °C, 5% $CO_2$.

*Isolation and culture of mouse fibroblast-like synoviocytes* The knee joints of mice were exposed by removing the skin, the quadriceps muscles were then resected in the middle and pulled in a distal direction to expose the patella. Patellae were collected in PBS before transfer to FLS media: Dulbecco's Modified Eagle Medium/F-12 Nutrient Mixture + GlutaMAX (Life Technologies) supplemented with 25% (v/v) FBS and 1% (v/v) penicillin/streptomycin in a 24-well plate and incubated at 37 °C, 5% $CO_2$. Media was refreshed daily. Once a substantial number of FLS had grown out of the patellae, the tissue was discarded and remaining adherent cells trypsinised, cells from two patellae were combined in 6-well plates and maintained by replacing culture media every other day. FLS were used for experiments from passage 3 (P3), as from this point FLS markers have been shown to be enriched, with low endothelial and macrophage marker expression[2]. Once confluent, P3 FLS were serum-starved for 6-hours before stimulation with either 100 μM BTB or an equivalent concentration of DMSO prepared in unsupplemented Dulbecco's Modified Eagle Medium/F12 Nutrient Mixture + GlutaMAX; a control condition of cells incubated solely in unsupplemented media was included for each experiment. 24-hours post-stimulation, culture media was collected and stored at -80 °C.

*Isolation and culture of human fibroblast-like synoviocytes* FLS were isolated from joint synovial tissues collected peri-operatively from consenting OA patients following elective total joint replacement under ethical approval granted by UK National Research Ethics Committee (NRES 16/SS/0172). Synovial tissue was diced (~1mm) and plated in T75 tissue culture flasks, incubated at 37 °C, 5% CO2 in complete fibroblast growth media (RPMI-1640 media (Sigma-Aldrich) supplemented with 10% foetal calf serum (Sigma-Aldrich), 1% non-essential amino acids (Sigma-Aldrich), 1% sodium orthopyruvate (Sigma-Aldrich), 2 mM L-glutamine (ThermoFisher Scientific), 1% penicillin and streptomycin (100 U/mL penicillin and 100 μg/mL streptomycin, Sigma-Aldrich). Once fibroblasts had migrated out of tissue, tissues were discarded and adherent cells were maintained in culture till ~70% confluent, with media changes every 3-4 days. FLS used in experiments were from passage 4 (P4) which we have previously shown are GP38 positive fibroblasts[3]. ~200,000 cells were plated per well of a 6-well plate. Prior to treatments, media was changed to 0% FSC media overnight. The following day cells were stimulated with 100 μM BTB or an equivalent concentration of DMSO prepared in 0% FSC media, 24-hours later culture media was collected and stored at -80 °C.

*Molecular cloning* Phusion high-fidelity DNA polymerase (Thermo Fisher) was used to amplify the open reading frame of genes of interest from mouse cDNA libraries prepared from RNA extracted from an appropriate tissue (reported to express the gene of interest). Primers were designed to introduce flanking restriction sites and N- and C-terminal tags as necessary to permit subcloning into a variety of recipient plasmids following standard restriction cloning practices. The pcDNA5/FRT/TO vector was purchased from Thermo Fisher. A construct encoding protease-activated receptor-2 tagged with Renilla luciferase 8 (RLuc8), a kind gift from Prof. Nigel Bunnett (New York University) was used to create GPR65-RLuc8 via subcloning. All constructs were verified by Sanger sequencing (Eurofins Genomics).

*Cell line culture* All cell lines were cultured at 37 °C in a humidified 5% $CO_2$ incubator. Flp-In CHO cells (Thermo Fisher) were maintained in F12 media supplemented with 10% (v/v) FBS, 1% (v/v) penicillin/streptomycin and 100 μg/ml zeocin (InvivoGen). Cells stably expressing PS-GPCR constructs were generated by co-transfecting Flp-In CHO cells with the desired PS-GPCR construct (cloned into a pcDNA5/FRT/TO vector) and the Flp recombinase, pOG44, at a ratio of 1:9, PS-GPCR:pOG44 using lipofectamine LTX and in the absence of any antibiotics. 48-hours post-transfection, culture media was replaced with standard CHO media containing 550 µg/ml hygromycin-B (InvivoGen). After expansion of hygromycin-B resistant cells, expression of constructs was confirmed using reverse transcription PCR (RT-PCR) and immunocytochemistry.

*qPCR* RNA was isolated from FLS using TRIzol (Merck): following trypsinisation cells were resuspended in 700 µl TRIzol and passed through a 21G needle several times, after a 5-min incubation 100 µl 1-bromo-3-chloropropane was added and samples were shaken vigorously before a further 3-minute incubation. Samples were then centrifuged (12,000g, 15-minutes) at 4 °C to separate the different phases, the least dense, colourless phase, containing RNA, was collected and cleaned up using the spin columns and wash buffers supplied with Zymo Research kits. High-capacity reverse transcription reagents (Applied Biosystems) were used to synthesise complementary DNA (cDNA) from RNA samples using random hexamers to prime reactions. Reverse transcription reactions included 500 ng of RNA as a template, the efficiency of reverse transcription was assumed to be 100% (i.e. 500 ng RNA assumed to produce 500 ng cDNA). Controls where reverse transcriptase was omitted from the reaction mix were performed in parallel. cDNA samples were diluted to 2.5 ng/µl in $ddH_2O$ and gene expression was assessed using TaqMan assays (Thermo Fisher) as per the manufacturer's guidelines. Thermocycling was performed using a StepOnePlusTM System (Thermo Fisher) and the following settings: 50 °C for 2-minutes, 95 °C for 10-minutes, 40 cycles of 95 °C for 15-seconds and 60 °C for 1-minute. The fluorescence intensity of samples was captured during the final minute of each cycle. Each sample of cDNA was assayed in duplicate per gene of interest, alongside appropriate negative controls where a combination of RNA samples was used as the template. The probes used were: Gpr4 (Mm00558777_s1), Gpr31 (Mm02391728_g1), Gpr65 (Mm00433695_m1), Gpr68 (Mm01335275_m1), Gpr132 (Mm00490809_m1), Gpr151 (Mm00808987_s1), Cdh-11 (Mm00515466_m1), Cd-248 (Mm00547485_s1), Cd-68 (Mm03047343_m1), Cd-31 (Mm01242576_m1) and 18SrRNA (Mm03928990_g1). Relative expression was calculated as $2^{-\Delta Ct}$, where $\Delta$Ct is the difference in the Ct value obtained for the gene of interest minus that of the housekeeping gene, *18SrRNA.*

*Immunostaining* FLS were fixed by exposure to ice-cold 100% methanol for 5-minutes at -20 °C, after which cells were washed with PBS three times. Cells were incubated at room temperature with antibody dilutant (containing donkey serum and BSA) for an hour before overnight incubation with primary antibodies at 4 °C. Primary antibodies used were αCDH-11 (1:100, rabbit polyclonal, Invitrogen, 71-7600) and αpCREB (1:500, rabbit polyclonal, Cell Signalling Technology, 9198). The following day FLS were washed with PBS containing 0.001% (v/v) Tween-20 three times before incubation with arabbit Ig-AF488 (1:1000, donkey polyclonal, Invitrogen, A32790) for 2-hours at room temperature, followed by a further two washes with PBS-Tween20. The nucleus was stained with DAPI for 2-minutes, after two washes, cells were left in PBS and imaged with a Leica SP5 laser-scanning confocal microscope and 63x oil objective. The intensity of pCREB staining was analysed in ImageJ, briefly, cells were selected as regions of interest on the brightfield image and intensity of 488 channel was then measured. The intensity for a random area of background was subtracted and intensities were normalised to the total area of the cell.

*Antibody array to assess inflammatory mediator release by FLS cells* Collected conditioned FLS media were analysed with Mouse or Human Inflammation Antibody Array (Abcam) as appropriate, following the manufacturer's guidelines. A total of 2 ml conditioned media, pooled from three (mouse FLS), or four (human FLS) separate biological replicates, was incubated with membranes overnight at 4 °C. Chemiluminescent signals were captured with an Odyssey XF imager (Li-Cor). Densitometry analysis was performed in ImageJ: for each array the intensities of targets were normalised to the average intensity of positive control spots, yielding relative density. Fold-change in concentrations were calculated as the relative density for test agonists – vehicle control / vehicle control, analyte-by-analyte.

*BRET assays* Flp-In CHO cells were plated in 100 mm dishes, once 80% confluent, cells were transiently transfected with 4 µg β-arrestin1-YFP/β-arrestin2-YFP/RIT-Venus/pcDNA3.1 and 1 µg mGPR65-RLuc8 using Lipofectamine LTX transfection reagent (3:1, LTX:cDNA). Fluorescently tagged BRET acceptor constructs were a kind gift from Prof. Nigel Bunnett (New York University). 24-hours post-transfection, cells were resuspended in media and seeded in poly-D-lysine coated white 96-well OptiPlates (PerkinElmer) at a density of 30,000 cells / well. The following day, culture media was replaced with 100 µl HBSS containing 10 mM HEPES, 10 mM MES and 2 mg/ml BSA, pH 7.40. Cells were allowed to acclimatise for 30 minutes at 37 °C before the HBSS was replaced with 90 µl stimulatory solutions containing various concentrations of agonist or pH buffered as necessary. Cells were incubated (37 °C) in stimulatory solutions for 10-minutes before the addition of 5 µM coelenterazine-H. After a further 5-minutes incubation the intensities of RLuc8 and YFP luminescence were measured using a CLARIOstar plate reader (BMG LabTech) with bandwidth settings 475 ± 30 nm and 535 ± 30 nm respectively, each stimulation was assayed in duplicate. BRET ratios were calculated by dividing YFP emission by RLuc8 luminescence, ratios from cells expressing mGPR65-RLuc8 and pcDNA3.1 were subtracted from those expressing -YFP/-Venus constructs to yield netBRET ratios.

*cAMP assays (Cell lines)* The day before experiments cells were seeded in standard 96-well plates at a density of 15,000 cells / well, the following day culture media was replaced with HBBS containing 10 mM HEPES, 10 mM MES and 2 mg/ml BSA, pH 7.40. Cells were allowed to acclimatise for 30-minutes at 37 °C before the HBSS was replaced with 100 µl solutions of HBSS containing various concentrations of agonist or pH buffered as necessary, for testing psychosine solutions were also prepared with 1 µM forskolin. After 15 minutes of stimulation at 37 °C, plates were transferred to ice, stimulatory solutions were aspirated and 50 µl lysis buffer (50 mM HEPES, 10 mM $CaCl_2$, 0.35% Triton X-100; pH 7.40) was added to each well, the plate was incubated on ice with gentle agitation for 30-minutes to ensure full cells lysis. Each condition was assayed in duplicate. The cAMP in lysates was quantified using LANCE Ultra cAMP detection reagents (Perkin Elmer). Briefly 10 µl of cell lysate was incubated with europium (Eu) labelled cAMP tracer and cAMP-specific antibody labelled with ULight™ dye in white ProxiPlate 384-well plates (Perkin Elmer) for 1 hour in reduced lighting as per the manufacturer's guidelines. After the incubation, TR-FRET was measured using a Mithras LB 940 microplate reader (Berthold Technologies) with excitation at 340 nm and emission recorded at 665 nm. TR-FRET signals were normalised to those achieved from cells stimulated with 10 µM forskolin.

*cAMP assays (FLS)* P3 FLS were plated in standard 24-well plates, once confluent media was replaced with HBSS containing 10 mM HEPES, 10 mM MES and 2 mg/ml BSA, pH 7.40. Cells were allowed to acclimatise for 30-minutes at 37 °C before the HBSS was replaced with 200 µl solutions of HBSS containing various concentrations of agonist or pH buffered as necessary a well stimulated with 100 µM forskolin was included in each experiment as a positive control. After 15-minutes of stimulation at 37 °C, plates were transferred to ice, stimulatory solutions were aspirated and 50 µl lysis buffer (50 mM HEPES, 10 mM $CaCl_2$, 0.35% Triton X-100; pH 7.40) was added to each well, the plate was incubated on ice with gentle agitation for 30-minutes to ensure full cells lysis. The cAMP in lysates was quantified using LANCE Ultra cAMP detection reagents (Perkin Elmer) as above, each lysate was assayed in duplicate alongside a cAMP standard curve, used to interpolate cAMP concentration of each lysate. To account for differences in cell number between conditions the total protein content of lysates was also assayed using Bradford reagent (Merck) and a standard curve made up of known amounts of BSA.

*$Ca^{2+}$ mobilisation assays* The day before assays were conducted, cells were seeded in black 96-well ViewPlates (PerkinElmer) at a density of 50,000 cells / well. The following day, cells were loaded with the $Ca^{2+}$-sensitive dye Fluo4 (Thermo Fisher) by exchange of culture media for HBSS containing 10 mM HEPES, 10 mM MES, 2 mg/ml BSA, 250 mM probenecid and 2.5 µM Fluo4-AM. Cells were incubated for 45-minutes at 37 °C to ensure good uptake of the dye, after which cells were washed with HBSS containing 10 mM HEPES, 10 mM MES, 2 mg/ml BSA and 250 mM probenecid twice, cells were left in 90 µl of this assay buffer. Experiments were performed using a CLARIOstar microplate reader (BMG LabTech) using the inbuilt incubator set at 37 °C. The intensity of Fluo4 was monitored every second through excitation at 483 ± 14 nm and emission recorded at 530 ± 30 nm. Each well was monitored individually, after recording a 15 second baseline, the automated injectors of the microplate reader were used to dispense 10 µl of stimulatory solutions and measurements continued for a further 2-minutes, each stimulation was performed in duplicate. Stimulation-induced changes in $Ca^{2+}$mobilisation were observed after correcting Fluo4 intensity for the 15 second baseline of each recording, signals were then normalised to an Fmax achieved by stimulating cells with 10 µM ionomycin to yield F/Fmax, the peak values of which for each stimulation were used to plot concentration-response curves. Stimulatory solutions containing agonists were made at a 10 X concentration, to permit the desired concentration once injected to the 90 µl assay buffer. For proton-stimulation in $Ca^{2+}$ experiments, the concentration of 1 ml HCl required to change the pH of 9 ml pH 7.40 assay buffer to the desired pH was determined empirically before each experiment, the same change in pH was assumed to take place following injection of 10 µl HCl at the desired concentration to 90 µl assay buffer pH 7.40.

*ERK phosphorylation assays* Cells were plated at a density of 50,000 per well in standard 96-well plates, the following day culture media was replaced with serum-free Opti-MEM (Life Technologies). 16-hours later, cells were acclimatised in HBSS containing 10 mM HEPES, 10 mM MES, 2 mg/ml BSA, pH 7.40 for 30 minutes at 37 °C, after which assay buffer was exchanged for 100 µl stimulatory solutions (assay buffer containing agonists or buffered to a certain pH), cells were stimulated for 5 minutes at 37 °C before being placed on ice and lysed (50 mM HEPES, 10 mM $CaCl_2$, 0.35% Triton X-100; pH 7.40), plates were gently agitated for 30-minutes to ensure full cells lysis. All stimulations were performed in duplicate. Phosphorylated ERK1/2 was quantified in lysates using Phosphorylated ERK1/2 (T202-Y204) LANCE Ultra detection reagents. 15 µl of each lysate was combined with 0.5 nM Eu-labelledanti-ERK1/2 (T202/Y204) antibody and 5 nM ULight-labelled-anti-ERK1/2 antibody in white ProxiPlate 384-well plates in reduced lighting. After 4 hours of incubation TR-FRET was measured with a Mithras Plate Reader with excitation at 340 nm and emission recorded at 615 nm and 665 nm. Emission at 665 nm was divided by emission at 615 nm and data normalised to the ERK activation achieved by 10 µM PDBu.

*DRG neuron electrophysiology* Electrophysiology recordings were performed the day after DRG neuron cultures were prepared, using an EPC-10 amplifier (HEKA) and corresponding Patchmaster software. The extracellular solution contained (in mM): NaCl (140), KCl (4), $MgCl_2$ (1), $CaCl_2$ (2), glucose (4) and HEPES (10), pH 7.40. Patch pipettes were pulled from borosilicate glass capillaries (Hilgenberg) using a P-97 pipette puller (Sutter Instruments) with resistances of 4-8 MΩ and back-filled with intracellular solution containing (in mM): KCl (110), NaCl (10), $MgCl_2$ (1), EGTA (1), HEPES (10), $Na_2ATP$ (2), $Na_2GTP$ (0.5), pH 7.30. Whole cell currents or voltages were acquired at 20 kHz. Where necessary FB labelled neurons, identified by LED excitation at 365 nm (Cairn Research). Step-wise depolarisation (Δ10 pA, 50 ms) was used to determine the action potential threshold (rheobase) of cells. Only cells which fired action potentials were included in analyses. Action potential parameters were measured using Fitmaster software (HEKA) and IgorPro software (Wavemetrics). The excitability of neurons was further assessed by applying a suprathreshold (2x action potential threshold) for 500 ms, the frequency of action potentials during this time was noted. The activity of macroscopic voltage-gated channels was assessed in voltage-clamp mode with appropriate series resistance compensation. Cells were held at -120 mV for

150 ms before stepping to the test potential (-60 mV – 55 mV in 5 mV increments) for 40 ms and returning to a holding potential of -60 mV for 200 ms between steps. Peak inward and outward currents were normalised to cell size by dividing by cell capacitance. Peak inward current densities were then fit to a Boltzmann function to determine the reversal potential and half-activating potential of voltage-gated channels.

*Data analysis and statistics* Data are presented as mean ± standard error of the mean, the number of biological and technical replicates are detailed in individual figure legends. Agonist concentration response curves were fitted to a Hill equation, $y = min + ((max-min)/1+exp(Hill\ Coefficient*(EC_{50}-x)))$, empirically in R where *min* and *max* are the minimum and maximum response measured respectively, $EC_{50}$ is the concentration of agonist required to generate a response halfway between the *min* and *max*. To compare agonist profiles, the average maximal response of each agonist, in each pathway was normalised to that achieved by proton-stimulation (maximal responses were elicited by pH 5.8 for all pathways but cAMP accumulation, which peaked at pH 6.4). As psychosine inhibits forskolin-induced cAMP accumulation, the $E_{max}$ of psychosine in the cAMP pathway was set as 0. Appropriate analyses were selected according to the number of factors being compared and whether data met the assumptions for parametric analyses, the statistical tests employed are stated in corresponding figure legends. All analyses were performed in R, and $p$ values < 0.05 considered significant.

--


1. Radu, C. G. *et al.* Normal Immune Development and Glucocorticoid-Induced Thymocyte Apoptosis in Mice Deficient for the T-Cell Death-Associated Gene 8 Receptor. *Mol. Cell. Biol.* **26**, 668–677 (2006).

2. Chakrabarti, S. *et al.* Sensitization of knee-innervating sensory neurons by tumor necrosis factor-α-activated fibroblast-like synoviocytes: an in vitro, coculture model of inflammatory pain. *Pain* **161**, 2129–2141 (2020).

3. Farah, H. *et al.* Differential Metabotypes in Synovial Fibroblasts and Synovial Fluid in Hip Osteoarthritis Patients Support Inflammatory Responses. *Int. J. Mol. Sci.* **23**, 3266 (2022).

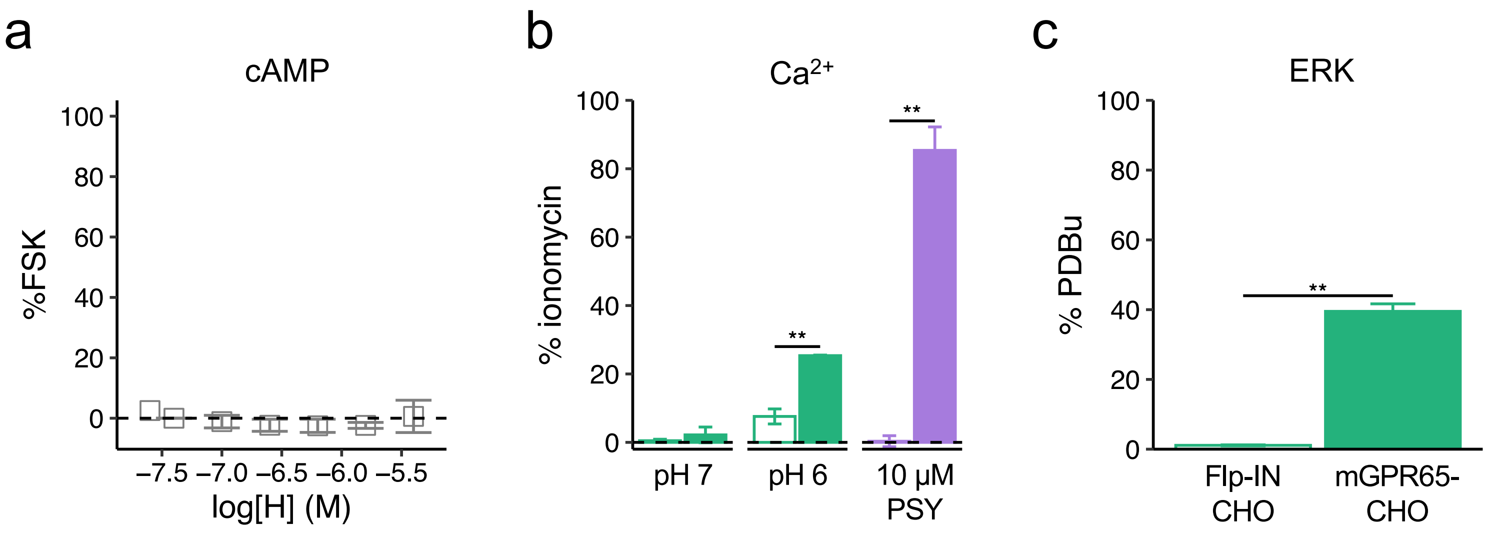


**Supplementary Figure 1. Assessing the GPR65 dependency of observed signalling in CHO cells. (a)** Stimulation of parental Flp-IN CHO cells with protons does not coordinate accumulation of cAMP. **(b)** Comparison of proton- and psychosine (PSY)-induced intracellular $Ca^{2+}$ mobilisation for Flp-IN CHO (open bars) and mGPR65-CHO cells (solid bars). **(c)** Comparison of proton-induced (pH 6) ERK activation for Flp-IN CHO (open bars) and mGPR65-CHO cells (solid bars). ** *p / p-adj* < 0.01: **(b)** Two-way ANOVA followed by Bonferroni-corrected post-hoc. (c) Unpaired t-test. Data are from at least two independent experiments where each [agonist] was assayed in duplicate.

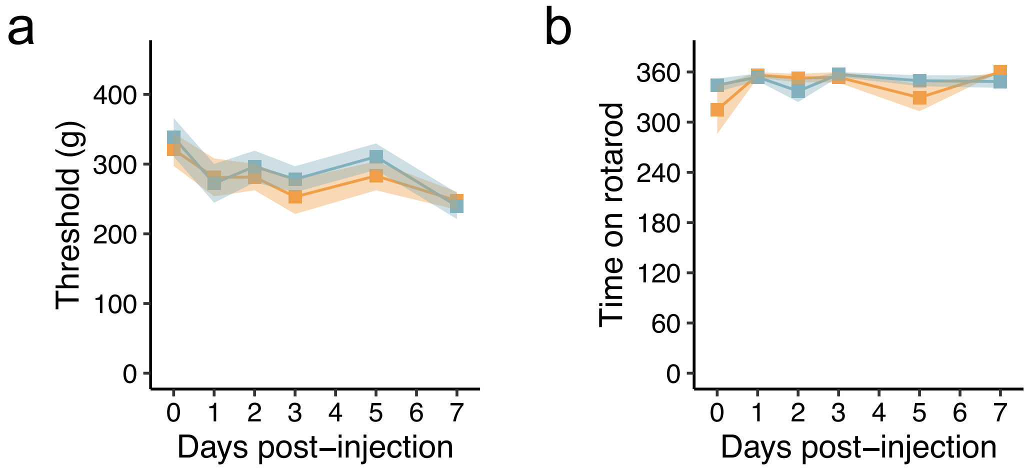


**Supplementary Figure 2. Mechanical threshold of the contralateral knee and locomotor function are not affected by intra-articular injection of BTB. (a)** The mechanical threshold of the non-injected knee joint was assessed by pressure application measurement. **(b)** Average time on rotarod across experimental timeline. N = 8 female mice per group.

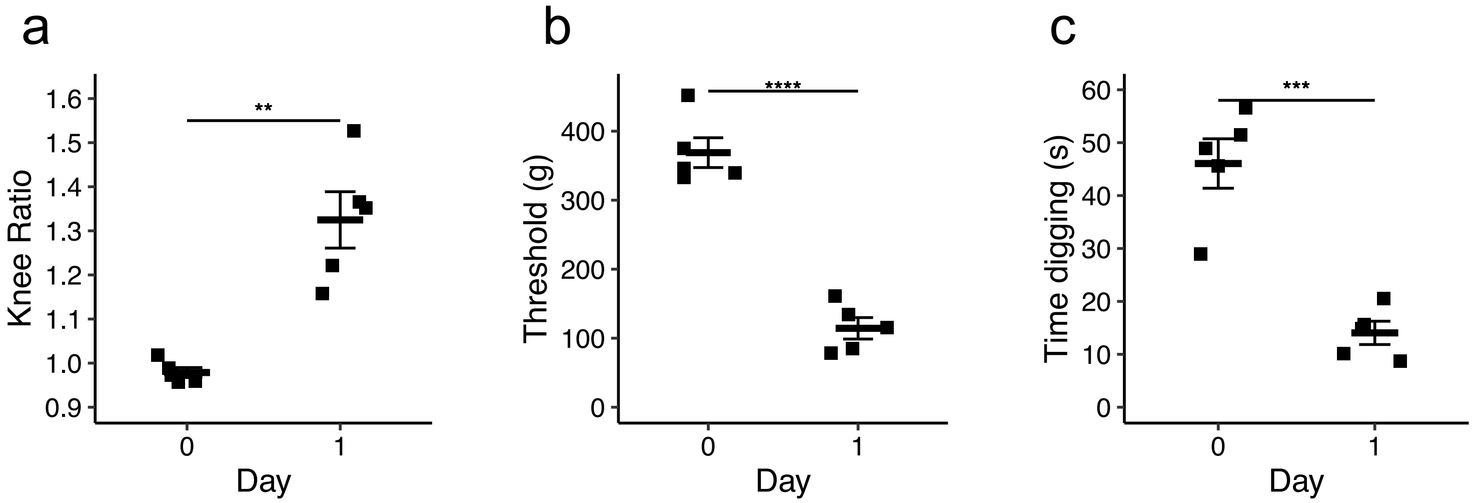


**Supplementary Figure 3. Inflammation and pain-like behaviours of wild-type mice following intra-articular injection of BTB before electrophysiological characterisation of knee-innervating neurons. (a)** The ratio of the ipsilateral to contralateral knee was calculated as a measure of the extent of inflammation. **(b)** Mechanical sensitivity of injected knee joints and **(c)** time spent digging were used to infer the pain status of mice. ** *p-adj* < 0.01, *** *p-adj* < 0.001: paired t-test. N = 5 mice.

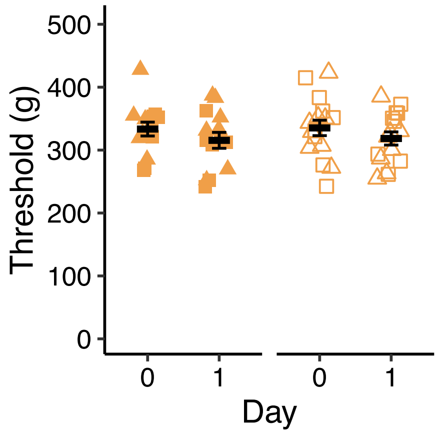


**Supplementary Figure 4. Mechanical threshold of the contralateral knee following intra-articular injection of BTB to WT and GPR65 KO mice. (a)** The mechanical threshold of the non-injected knee joint was assessed by pressure application measurement. N = 14 wild-type mice (7 female, 7 male) and 16 KO mice (8 female, 8 male).

**Supplemental Tables**

| Cytokine | group1 | group2 | p.adj | p.adj.signif |
|---|---|---|---|---|
| BLC | BTB | Control | 0.0582 | ns |
| BLC | BTB | DMSO | 0.143 | ns |
| BLC | Control | DMSO | 0.809 | ns |
| CD30L | BTB | Control | 0.0674 | ns |
| CD30L | BTB | DMSO | 0.0433 | * |
| CD30L | Control | DMSO | 1 | ns |
| Eotaxin | BTB | Control | 0.655 | ns |
| Eotaxin | BTB | DMSO | 0.135 | ns |
| Eotaxin | Control | DMSO | 0.527 | ns |
| Eotaxin-2 | BTB | Control | 0.277 | ns |
| Eotaxin-2 | BTB | DMSO | 0.0509 | ns |
| Eotaxin-2 | Control | DMSO | 0.292 | ns |
| Fas Ligand | BTB | Control | 1 | ns |
| Fas Ligand | BTB | DMSO | 0.704 | ns |
| Fas Ligand | Control | DMSO | 1 | ns |
| Fractalkine | BTB | Control | 0.797 | ns |
| Fractalkine | BTB | DMSO | 1 | ns |
| Fractalkine | Control | DMSO | 0.992 | ns |
| GCSF | BTB | Control | 0.352 | ns |
| GCSF | BTB | DMSO | 0.904 | ns |
| GCSF | Control | DMSO | 1 | ns |
| GM-CSF | BTB | Control | 0.929 | ns |
| GM-CSF | BTB | DMSO | 1 | ns |
| GM-CSF | Control | DMSO | 1 | ns |
| I-TAC | BTB | Control | 1 | ns |
| I-TAC | BTB | DMSO | 1 | ns |
| I-TAC | Control | DMSO | 1 | ns |
| IFNy | BTB | Control | 1 | ns |
| IFNy | BTB | DMSO | 1 | ns |
| IFNy | Control | DMSO | 1 | ns |
| IL-10 | BTB | Control | 0.155 | ns |
| IL-10 | BTB | DMSO | 0.29 | ns |
| IL-10 | Control | DMSO | 1 | ns |
| IL-12 p40/p70 | BTB | Control | 0.16 | ns |
| IL-12 p40/p70 | BTB | DMSO | 0.19 | ns |
| IL-12 p40/p70 | Control | DMSO | 1 | ns |
| IL-12 p70 | BTB | Control | 1 | ns |
| IL-12 p70 | BTB | DMSO | 0.93 | ns |
| IL-12 p70 | Control | DMSO | 1 | ns |
| IL-13 | BTB | Control | 1 | ns |

| | | | | |
|---|---|---|---|---|
| IL-13 | BTB | DMSO | 1 | ns |
| IL-13 | Control | DMSO | 1 | ns |
| IL-17 | BTB | Control | 0.522 | ns |
| IL-17 | BTB | DMSO | 1 | ns |
| IL-17 | Control | DMSO | 1 | ns |
| IL-1$\alpha$ | BTB | Control | 0.217 | ns |
| IL-1$\alpha$ | BTB | DMSO | 0.663 | ns |
| IL-1$\alpha$ | Control | DMSO | 0.968 | ns |
| IL-1$\beta$ | BTB | Control | 1 | ns |
| IL-1$\beta$ | BTB | DMSO | 0.999 | ns |
| IL-1$\beta$ | Control | DMSO | 1 | ns |
| IL-2 | BTB | Control | 1 | ns |
| IL-2 | BTB | DMSO | 1 | ns |
| IL-2 | Control | DMSO | 1 | ns |
| IL-4 | BTB | Control | 1 | ns |
| IL-4 | BTB | DMSO | 1 | ns |
| IL-4 | Control | DMSO | 1 | ns |
| IL-6 | BTB | Control | 1 | ns |
| IL-6 | BTB | DMSO | 1 | ns |
| IL-6 | Control | DMSO | 1 | ns |
| IL-7 | BTB | Control | 0.62 | ns |
| IL-7 | BTB | DMSO | 0.895 | ns |
| IL-7 | Control | DMSO | 1 | ns |
| IL-9 | BTB | Control | 0.491 | ns |
| IL-9 | BTB | DMSO | 0.786 | ns |
| IL-9 | Control | DMSO | 1 | ns |
| KC | BTB | Control | 0.472 | ns |
| KC | BTB | DMSO | 1 | ns |
| KC | Control | DMSO | 1 | ns |
| Leptin | BTB | Control | 0.0533 | ns |
| Leptin | BTB | DMSO | 0.43 | ns |
| Leptin | Control | DMSO | 0.208 | ns |
| LIX | BTB | Control | 0.0333 | * |
| LIX | BTB | DMSO | 0.17 | ns |
| LIX | Control | DMSO | 0.24 | ns |
| Lymphotactin | BTB | Control | 0.248 | ns |
| Lymphotactin | BTB | DMSO | 0.266 | ns |
| Lymphotactin | Control | DMSO | 1 | ns |
| MCP-1 | BTB | Control | 0.0382 | * |
| MCP-1 | BTB | DMSO | 0.00676 | ** |
| MCP-1 | Control | DMSO | 0.0633 | ns |
| MCSF | BTB | Control | 1 | ns |
| MCSF | BTB | DMSO | 1 | ns |
| MCSF | Control | DMSO | 1 | ns |

| | | | | |
|---|---|---|---|---|
| MIG | BTB | Control | 0.0807 | ns |
| MIG | BTB | DMSO | 0.0385 | * |
| MIG | Control | DMSO | 0.875 | ns |
| MIP-1α | BTB | Control | 0.31 | ns |
| MIP-1α | BTB | DMSO | 0.22 | ns |
| MIP-1α | Control | DMSO | 1 | ns |
| MIP-1γ | BTB | Control | 0.0073 | ** |
| MIP-1 γ | BTB | DMSO | 0.00437 | ** |
| MIP-1 γ | Control | DMSO | 0.496 | ns |
| RANTES | BTB | Control | 1 | ns |
| RANTES | BTB | DMSO | 1 | ns |
| RANTES | Control | DMSO | 1 | ns |
| SDF-1 | BTB | Control | 0.638 | ns |
| SDF-1 | BTB | DMSO | 0.739 | ns |
| SDF-1 | Control | DMSO | 1 | ns |
| sTNF RI | BTB | Control | 0.349 | ns |
| sTNF RI | BTB | DMSO | 0.0881 | ns |
| sTNF RI | Control | DMSO | 0.541 | ns |
| sTNF RII | BTB | Control | 1 | ns |
| sTNF RII | BTB | DMSO | 0.844 | ns |
| sTNF RII | Control | DMSO | 1 | ns |
| TCA-3 | BTB | Control | 0.744 | ns |
| TCA-3 | BTB | DMSO | 0.613 | ns |
| TCA-3 | Control | DMSO | 0.167 | ns |
| TECK | BTB | Control | 0.527 | ns |
| TECK | BTB | DMSO | 1 | ns |
| TECK | Control | DMSO | 0.41 | ns |
| TIMP-1 | BTB | Control | 1 | ns |
| TIMP-1 | BTB | DMSO | 0.777 | ns |
| TIMP-1 | Control | DMSO | 0.782 | ns |
| TIMP-2 | BTB | Control | 0.336 | ns |
| TIMP-2 | BTB | DMSO | 0.3 | ns |
| TIMP-2 | Control | DMSO | 1 | ns |
| TNF-α | BTB | Control | 0.759 | ns |
| TNF-α | BTB | DMSO | 0.418 | ns |
| TNF-α | Control | DMSO | 1 | ns |

Table S1 Comparison of inflammatory cytokine levels in the conditioned media collected from mouse FLS after 24 hours culture in regular media (Control), DMSO or BTB. * *p-adj* < 0.05, ** *p-adj* < 0.01: Two-way ANOVA followed by Bonferroni-corrected post-hoc.

| Cytokine | p.adj | p.adj.signif |
|---|---|---|
| BLC | 0.0156 | * |
| CD30 L | 0.00371 | ** |
| Fractalkine | 0.551 | ns |
| GCSF | 0.00409 | ** |
| IL-12 p70 | 0.0332 | * |
| IL-13 | 0.058 | ns |
| IL-1$\alpha$ | 0.359 | ns |
| IL-4 | 0.152 | ns |
| IL-6 | 0.00935 | ** |
| IL-9 | 0.0491 | * |
| KC | 0.00442 | ** |
| LIX | 0.0385 | * |
| Lymphotactin | 0.0433 | * |
| MCP-1 | 0.000713 | *** |
| MCSF | 0.0967 | ns |
| MIG | 0.0569 | ns |
| MIP-1$\alpha$ | 0.0172 | * |
| MIP-1 $\gamma$ | 0.00573 | ** |
| RANTES | 0.0103 | * |
| SDF-1 | 0.821 | ns |
| TCA-3 | 0.0551 | ns |
| TECK | 0.0402 | * |
| TIMP-1 | 0.455 | ns |
| TIMP-2 | 0.185 | ns |
| TNF-$\alpha$ | 0.0938 | ns |
| sTNF RI | 0.00923 | ** |
| sTNF RII | 0.0128 | * |

Table S2 Comparison of inflammatory cytokine levels in the conditioned media collected from WT versus KO mouse FLS after 24 hours stimulation with BTB. * *p-adj* < 0.05, ** *p-adj* < 0.01, *** *p-adj* < 0.001: Two-way ANOVA followed by Bonferroni-corrected post-hoc.

| Gene Symbol | Non-pain (Average TMM) | Pain (Average TMM) | p-value | q-value | Difference | Fold change |
|---|---|---|---|---|---|---|
| *TRPM7* | 2.4909 | 3.0907 | 0.64754306 | 0.85044065 | 0.59979999 | 1.51550645 |
| *GPR65* | 0.30077833 | 2.239871667 | 0.11665299 | 0.85044065 | 1.93909311 | 3.83464524 |
| *GPR4* | -0.732175 | -1.758886667 | 0.51986347 | 0.85044065 | -1.0267117 | 0.49082761 |
| *KCNK3* | -0.8157217 | -1.199538333 | 0.83165701 | 0.87919312 | -0.3838167 | 0.76640732 |
| *KCNK2* | -1.06275 | 2.419266667 | 0.003587 ** | 0.85044065 | 3.48201656 | 11.1735565 |
| *KCNK16* | -1.9628267 | -1.773283333 | 0.79213653 | 0.85044065 | 0.18954331 | 1.14040266 |
| *GPR132* | -2.0168 | -1.950074833 | 0.95997326 | 0.97089825 | 0.06672508 | 1.04733653 |
| *GPR31* | -2.10945 | -1.9199 | 0.79212501 | 0.85044065 | 0.18954997 | 1.14040792 |
| *TRPV4* | -2.2458 | -2.52415 | 0.81613345 | 0.8668191 | -0.27835 | 0.82453348 |
| *GPR68* | -2.3358 | 0.279746667 | 0.07409095 | 0.85044065 | 2.61554646 | 6.12855293 |
| *ASIC3* | -2.5258733 | -2.33635 | 0.79215565 | 0.85044065 | 0.18952331 | 1.14038685 |
| *GPR151* | -2.6941717 | -2.504633333 | 0.79213838 | 0.85044065 | 0.18953837 | 1.14039876 |
| *KCNK5* | -2.9870333 | -1.833935 | 0.20024279 | 0.85044065 | 1.15309823 | 2.22390972 |
| *TRPC4* | -3.1396333 | -2.950088 | 0.79213225 | 0.85044065 | 0.18954535 | 1.14040427 |
| *ASIC1* | -3.3256833 | -3.400288333 | 0.91904699 | 0.94274908 | -0.074605 | 0.9496021 |
| *ASIC4* | -3.4598333 | -3.27029 | 0.79213397 | 0.85044065 | 0.18954328 | 1.14040263 |
| *TRPV1* | -3.5319333 | -2.549905 | 0.3298347 | 0.85044065 | 0.98202837 | 1.97524056 |
| *ASIC2* | -3.55215 | -3.362613333 | 0.79214041 | 0.85044065 | 0.18953668 | 1.14039742 |
| *TRPC5* | -3.6132333 | -2.487906667 | 0.31484028 | 0.85044065 | 1.12532651 | 2.18150913 |
| *TRPA1* | -3.7331333 | -3.54359 | 0.79213386 | 0.85044065 | 0.18954338 | 1.14040272 |
| *KCNK9* | -3.8296333 | -3.640093333 | 0.79213653 | 0.85044065 | 0.18954012 | 1.14040014 |

Table S3 Summary of expression of known proton-sensitive receptors and channels from bulk synovium sequencing data of painful and non-painful sites from end-stage osteoarthritis patients. Data previously published by Nanus *et al.* (2021). Genes are ordered by expression level at non-painful sites, *GPR65* is highlighted. ** $p < 0.01$: Published differentially expressed gene (DEG) data from Nanus *et al.* (2021) was filtered to identify DEGs using two-group statistical comparison for > 1.5 fold change and $p < 0.05$.

| Cytokine | p.adj | p.adj.signif |
|---|---|---|
| IFNy | 0.0684 | ns |
| IL-11 | 0.49 | ns |
| IL-15 | 0.653 | ns |
| IL-17 | 0.399 | ns |
| IL-1β | 0.603 | ns |
| IL-4 | 0.015 | * |
| IL-6 | 0.000678 | *** |
| IL-8 | 0.00107 | ** |
| IP10 | 0.258 | ns |
| M-CSF | 0.765 | ns |
| MCP-1 | 0.00119 | ** |
| MCP-2 | 0.431 | ns |
| MIG | 0.95 | ns |
| MIP-1α | 0.0707 | ns |
| MIP-1β | 0.725 | ns |
| PDGFBB | 0.389 | ns |
| RANTES | 0.108 | ns |
| TIMP2 | 0.0917 | ns |
| TNF-α | 0.458 | ns |

Table S4 Comparison of inflammatory cytokine levels in the conditioned media collected from FLS isolated from human OA patients, from painful sites, 24-hours post-stimulation with BTB. * *p-adj* < 0.05, ** *p-adj* < 0.01, *** *p-adj* < 0.001: Two-way ANOVA followed by Bonferroni-corrected post-hoc.

| Sample ID | Sex | Age | BMI | Current medication(s) |
|---|---|---|---|---|
| 009 | M | 82 | 34.18 | Lansoprazole, Amlodipine |
| 015 | M | 84 | 25.29 | Lansoprazole, Aspirin, Atorvastatin, Ilbestoton, Gliclazide, Latanoprost, Lercanidipine, Hiagliplin |
| 028 | M | 74 | 27.68 | Co-codamol |
| 030 | M | 68 | 27.07 | Co-codamol |
| 034 | F | 67 | 30.86 | Co-codamol |
| 062 | M | 63 | 32.53 | Paracetamol |

Table S5 Synovial fluid donor details.
